# Reprogramming human T cell function and specificity with non-viral genome targeting

**DOI:** 10.1101/183418

**Authors:** Theodore L. Roth, Cristina Puig-Saus, Ruby Yu, Eric Shifrut, Julia Carnevale, Joseph Hiatt, Justin Saco, Han Li, Jonathan Li, Victoria Tobin, David Nguyen, Andrea M. Ferris, Jeff Chen, Jean-Nicolas Schickel, Laurence Pellerin, David Carmody, Gorka Alkorta-Aranburu, Daniela Del Gaudio, Hiroyuki Matsumoto, Montse Morell, Ying Mao, Rolen Quadros, Channabasavaiah Gurumurthy, Baz Smith, Michael Haugwitz, Stephen H. Hughes, Jonathan Weissman, Kathrin Schumann, Andrew P. May, Alan Ashworth, Gary Kupfer, Siri Greeley, Rosa Bacchetta, Eric Meffre, Maria Grazia Roncarolo, Neil Romberg, Kevan C. Herold, Antoni Ribas, Manuel D. Leonetti, Alexander Marson

## Abstract

Human T cells are central to physiological immune homeostasis, which protects us from pathogens without collateral autoimmune inflammation. They are also the main effectors in most current cancer immunotherapy strategies^1^. Several decades of work have aimed to genetically reprogram T cells for therapeutic purposes^2–5^, but as human T cells are resistant to most standard methods of large DNA insertion these approaches have relied on recombinant viral vectors, which do not target transgenes to specific genomic sites^6, 7^. In addition, the need for viral vectors has slowed down research and clinical use as their manufacturing and testing is lengthy and expensive. Genome editing brought the promise of specific and efficient insertion of large transgenes into target cells through homology-directed repair (HDR), but to date in human T cells this still requires viral transduction^8, 9^. Here, we developed a non-viral, CRISPR-Cas9 genome targeting system that permits the rapid and efficient insertion of individual or multiplexed large (>1 kilobase) DNA sequences at specific sites in the genomes of primary human T cells while preserving cell viability and function. We successfully tested the potential therapeutic use of this approach in two settings. First, we corrected a pathogenic *IL2RA* mutation in primary T cells from multiple family members with monogenic autoimmune disease and demonstrated enhanced signalling function. Second, we replaced the endogenous T cell receptor (*TCR*) locus with a new TCR redirecting T cells to a cancer antigen. The resulting TCR-engineered T cells specifically recognized the tumour antigen, with concomitant cytokine release and tumour cell killing. Taken together, these studies provide preclinical evidence that non-viral genome targeting will enable rapid and flexible experimental manipulation and therapeutic engineering of primary human immune cells.

---

The major barrier to effective non-viral T cell genome targeting of large DNA sequences has been the toxicity of the DNA^10^. While the introduction of short single stranded oligodeoxynucleotide (ssODN) HDR templates does not cause significant T cell death, it has been shown that larger linear double stranded (dsDNA) templates are toxic at high concentrations^11, 12^. Contrary to expectations, we found that co-electroporation of human primary T cells with CRISPR-Cas9 ribonucleoprotein (Cas9 RNP^13, 14^) complexes and long (>1kb) linear dsDNA templates reduced the toxicity associated with the dsDNA template (**Extended Data Fig 1**). Cas9 RNPs were co-electroporated with a dsDNA HDR template designed to introduce an N-terminal GFP-fusion in the housekeeping gene *RAB11A* (**Fig. 1a**). Systematic exploration of this approach while optimizing for both viability and efficiency (**Fig. 1b** and **Extended Data Fig. 2**) resulted in GFP expression in ~50% of cells in both primary human CD4+ and CD8+ T cells. The method was reproducibly efficient while maintaining high cell viability and expandability (**Fig. 1c, d, e,** and **Extended Data Fig. 3**). The system is also compatible with current manufacturing protocols for cell therapies as it could be applied to fresh or cryopreserved cells, bulk T cells or FACS-sorted sub-populations, and cells from whole blood or leukapheresis (**Extended Data Fig. 4**).

**Figure 1:**
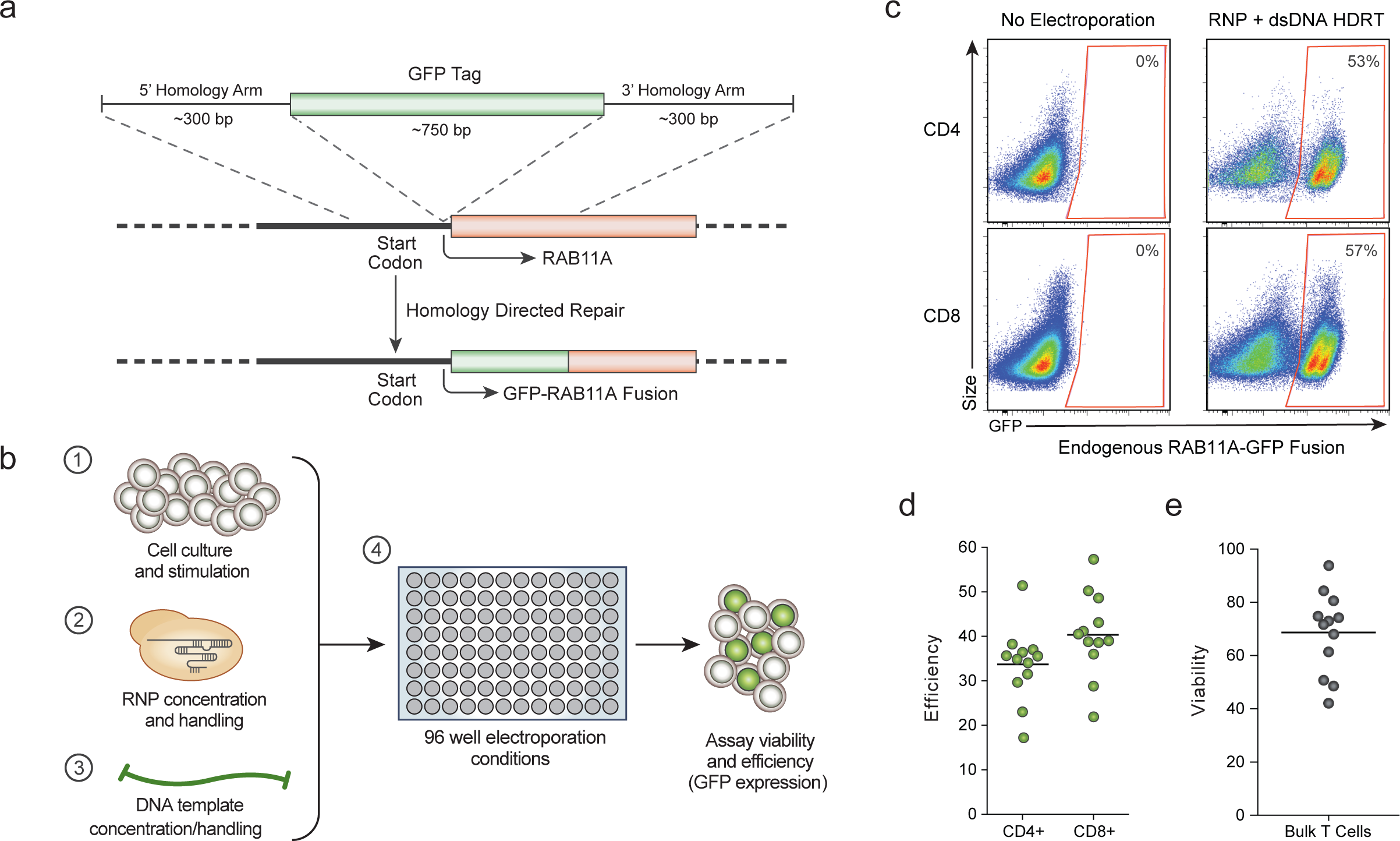
Efficient non-viral genome targeting in primary human T cells. **a,** Schematic diagram of a long (1350bp) linear dsDNA template encoding a GFP sequence flanked by regions homologous to the N-terminus of the housekeeping gene *RAB11A* (not drawn to scale). When a dsDNA break is induced at the N-terminus of *RAB11A,* the GFP sequence can be seamlessly introduced via homology directed repair (HDR) to generate an endogenously-tagged RAB11A-GFP fusion protein. **b,** Systematic analysis of the effects of cell culture and stimulation conditions, RNP and DNA template formulations, and electroporation conditions using 96-well high-throughput electroporations enabled rapid optimization of both cell viability (total number of live cells in culture) and HDR efficiency (% of cells GFP positive). **c,** High-efficiency insertion of a GFP fusion into the endogenous *RAB11A* gene was achieved using non-viral targeting in both primary human CD4+ and CD8+ T cells. **d,** Efficiency of targeted integration of the RAB11A-GFP HDR template in a cohort of 12 healthy human donors. Average efficiency was 33.7% and 40.3% in CD4+ and CD8+ primary human T cells respectively. **e,** Viability of primary human T cells after non-viral genome targeting relative to nonelectroporated controls averaged 68.6%. Efficiency and viability were measured 4 days following electroporation.

We next confirmed that the system could be applied broadly by targeting sequences in different locations throughout the genome. We efficiently engineered GFP+ primary T cells by generating fusions with different genes (**Fig. 2a** and **Extended Data Fig. 5**). Live-cell imaging with confocal microscopy confirmed the specificity of gene targeting, revealing the distinct sub-cellular locations of each resulting GFP-fusion protein^15^ (**Fig. 2b**). Appropriate chromatin binding of a transcription factor GFP-fusion protein was assessed by performing genome-wide CUT & RUN^16^ analysis with an anti-GFP antibody (**Fig. 2c** and **Extended Data Fig. 6**). Finally, we validated that gene targeting preserved endogenous gene regulation. Consistent with correct cell-type specific expression, a CD4-GFP fusion was selectively expressed in CD4+ T cells, but not in CD8+ T cells (**Fig. 2d**). Using HDR templates encoding multiple fluorescent proteins, we demonstrated that we could generate cells with bi-allelic gene targeting (**Fig. 2e** and **Extended Data Fig. 7**) or multiplex modification of two (**Fig. 2f** and **Extended Data Fig. 8**) or even three (**Fig. 2e** and **Extended Data Fig. 8**) different genes^17, 18^. Individual or multiple endogenous genes can therefore be directly engineered without virus in T cells, preserving gene and protein regulation.

**Figure 2:**
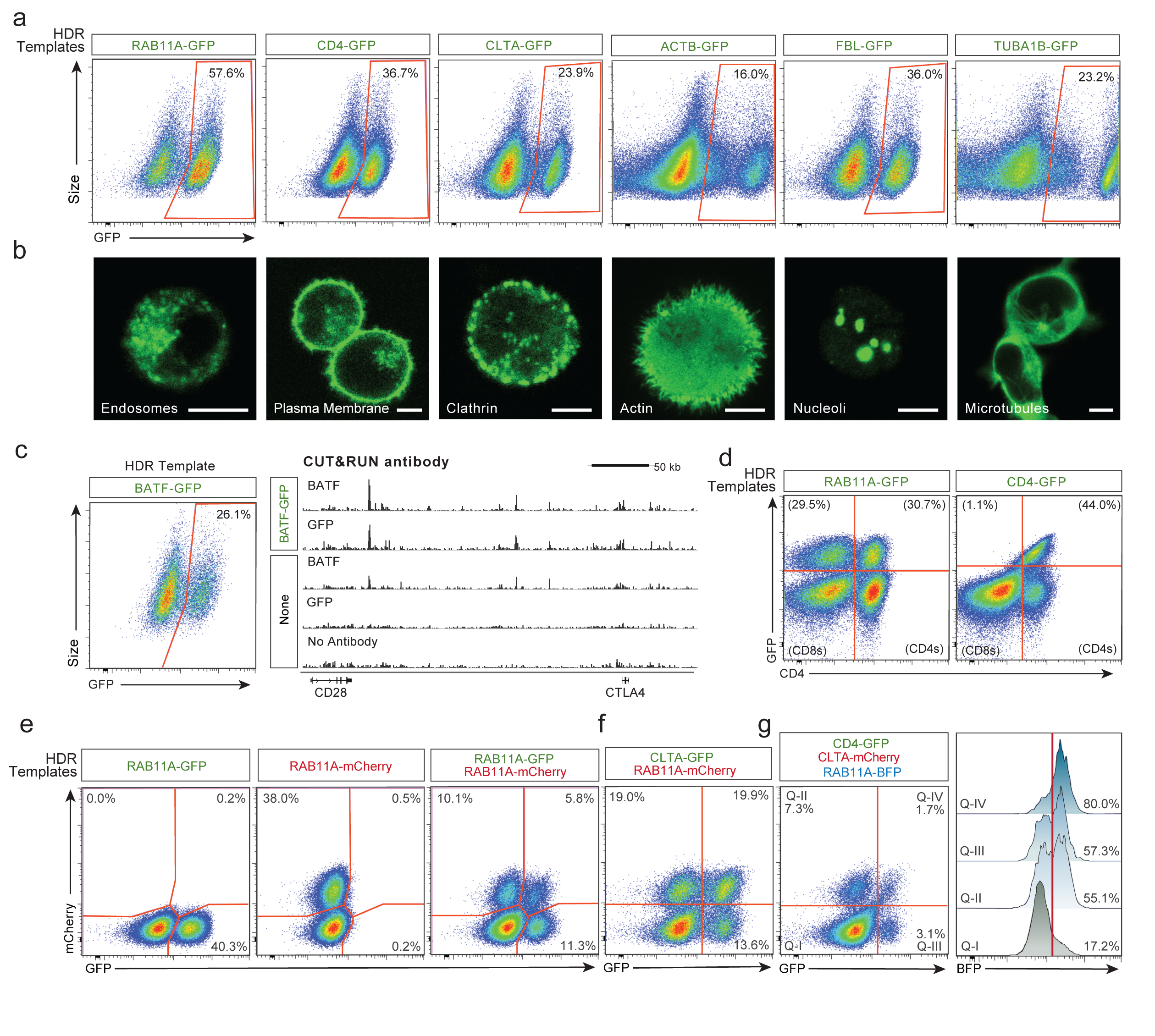
Individual and multiplexed modification of endogenous T cell genes. **a,** High efficiency genome targeting with GFP-fusion constructs could be achieved in multiple endogenous genes in primary human T cells using non-viral HDR templates and corresponding RNPs. **b,** Confocal microscopy of living, primary human T cells 7 days after electroporation of the indicated HDR template confirmed the appropriate localization of the fusion-protein. Scale bar in each image is 5 μm. Images are representative of two donors. **c,** Primary human T cells were engineered to express GFP fused to the endogenous transcription factor, BATF. At 11 Days post electroporation, nuclei were isolated and CUT&RUN was performed. GFP-BATF and total BATF chromatin interaction sites were identified using anti-GFP or anti-BATF antibodies. **d,** Non-viral targeting of GFP-fusion constructs to the *RAB11A* and *CD4* genes in bulk human primary T cells. RAB11A-fusions produced GFP positive CD4+ and CD8+ cells, whereas the CD4-fusions were selectively expressed in CD4+ cells. Note the linear relationship between GFP expression and CD4 staining consistent with specific fusion. **e,** Simultaneous electroporation of HDR templates to create RAB11A-GFP and/or RAB11A-mCherry fusions in primary human T cells. A distinct population of dual GFP+ mCherry+ cells was found when both templates are introduced concurrently, consistent with targeting both alleles. **f,** Multiplexed integration of HDR templates at two separate genomic loci was achieved in primary human T cells. **g,** Simultaneous targeting of three distinct genomic loci was successful. Cells positive for a single integration (Q-II, Q-III) and even further two integrations (Q-IV) were highly enriched for integration of the third HDR template. Flow cytometry was used to assay efficiency (performed at 4 days after electroporation).

For therapeutic use of genetically modified T cells, integrated sequences should be introduced specifically without unintended disruption of other critical genome sites^19^. We performed targeted locus amplification (TLA) sequencing^20^ and found no evidence of off-target integrations above the assay’s limit of detection (~1% of alleles) (**Extended Data Fig. 9**). We further assessed potential off-target integrations at the single cell level by quantifying GFP+ cells generated with a Cas9 RNP that cuts outside the homology site. Similar to what has been described with viral HDR templates^8, 21^, we found evidence to suggest that double-stranded templates could integrate independent of target homology^22, 23^, albeit at low rates (**Extended Data Fig. 10**). These rare events could be reduced almost completely by using single-stranded DNA templates^24, 25^ (**Extended Data Fig. 11**). As an additional safeguard that could be important for some applications, we demonstrated that non-viral T cell genome targeting also could be achieved using either a single-stranded or double-stranded template with a Cas9 “nickase” engineered to reduce potential off-target double-stranded cuts^26, 27^ (**Extended Data Fig. 12**).

Having optimized this non-viral gene engineering approach in primary human T cells, we demonstrated its utility it in two different clinically relevant settings where targeted replacement of a gene would provide proof-of-principle for therapeutic effect in patients. Specifically, we tested the ability to rapidly and efficiently correct an inherited T cell genetic alteration and we also tested the targeted insertion of the two chains of a TCR to genetically redirect the specificity of T cells to recognize cancer.

We identified a family with monogenic primary immune dysregulation with autoimmune disease caused by recessive loss-of-function mutations in the gene encoding the IL-2 alpha receptor (IL2RA)^28^(**Extended Data Table 1**), which is essential for healthy regulatory T cells (Tregs)^29, 30^ (**Extended Data Fig. 13**). Whole exome sequencing revealed that the IL2RA-deficient children harboured compound heterozygous mutations in *IL2RA* (**Fig. 3a** and **Extended Data Fig. 14**). One mutation, c.530A>G, creates a premature stop codon. With non-viral genome targeting, we were able to correct the mutation and observe IL2RA expression on the surface of corrected T cells from the patient (**Fig. 3b**). Long dsDNA templates led to efficient correction of the mutations. Because only two base pair changes were necessary (one to correct the mutation and one to silently remove the gRNA’s PAM sequence), a short single-stranded DNA (~120 bps) could also be used to make the correction. These single-stranded DNAs were able to correct the mutation at high frequencies, although the efficiency of correction was lower than with the longer dsDNA template (**Extended Data Fig. 15**, **16**). Correction was successful in T cells from all three siblings, but lower rates of IL2RA expression were seen in compound het 3, which could be due to altered cell-state associated with the patient’s disease or the fact she was the only sibling treated with immunosuppressive therapy (**Extended Data Table 1** and **Extended Data Fig. 17**). The second mutation identified, c.800delA, causes a frameshift in the reading frame of the final *IL2RA* exon. This frameshift mutation could be corrected both by HDR as well as by RNP cutting alone, presumably due to small indels restoring the reading frame (**Extended Data Fig. 16**). Taken together, these data show that distinct mutations can be corrected in patient T cells using HDR template-dependent and non-HDR template-dependent mechanisms.

**Figure 3:**
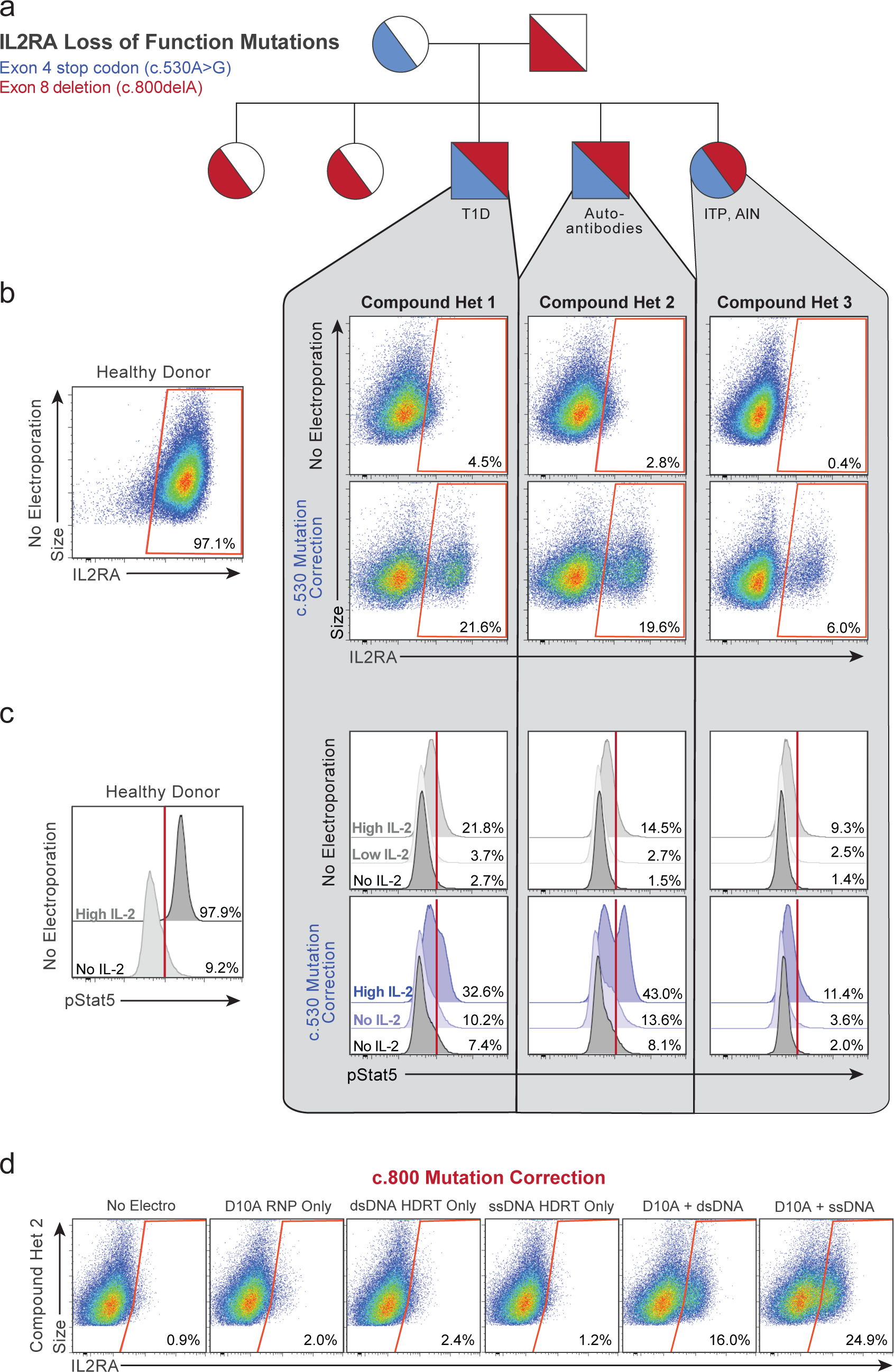
Monogenic autoimmune mutations corrected by non-viral genome targeting. **a,** Three siblings in a family carry two different *IL2RA* (encoding high-affinity IL-2 receptor, CD25) mutations (c.530A>G creating a stop codon in *IL2RA* exon 4; c.800delA, creating a frameshift mutation in *IL2RA* exon 8 which causes an almost 100 amino acid run-on). **b,** These three compound heterozygote siblings show greatly reduced, but not completely absent, cell surface expression of IL2RA on their primary T cells. Non-viral gene targeting of the c.530 mutation by electroporation of a Cas9 RNP and a dsDNA HDR template containing the correct *IL2RA* sequence (along with a silent mutation in the targeted PAM sequence) successfully rescued IL2RA cell surface expression in a portion of T cells from each compound heterozygote sibling 2 days following electroporation. **c,** 7 days after non-viral gene targeting, targeted T cells showed increased phosphorylation levels of Stat5 upon IL-2 stimulation compared to non-targeted controls. **d,** Non-viral gene targeting and correction of the second c.800 mutation was also efficient using D10A nickase along with a ssDNA HDR template. T cells were stained for IL2RA surface expression after 9 days of *ex-vivo* expansion following electroporation (2 days following re-stimulation).

Mutation correction improved cell signalling function. Following correction of the c.530A>G *IL2RA* mutation, IL-2 treatment led to increased STAT5 phosphorylation, a hallmark of productive signalling (**Fig. 3c** and **Extended Data Fig. 15**, **16**). In addition, following correction, the fraction of total T cells expressing both IL2RA and FOXP3, a critical transcriptional factor in Tregs, returned to levels similar to healthy controls (**Extended Data Fig. 15**, **16**). We were also able to correct IL2RA in a sorted population of CD3+CD4+CD127loTIGIT+CD45RO+ Treg-like patient cells (**Extended Data Fig. 15**), a potential strategy for a gene-modified cell therapy for the children in this family. Cell-type specific and stimulus responsive expression of *IL2RA* is under tight control by multiple endogenous c/s-regulatory elements that constitute a super-enhancer^31, 32^. Therefore, therapeutic correction of IL2RA is likely to depend on specifically repairing the gene in its endogenous genomic locus; off-target effects should be avoided. We therefore demonstrated that the c.800delA mutation could also be repaired using Cas9 nickase combined with a single-stranded HDR template (**Fig. 3d**).

Non-viral genome targeting not only allows the correction of point mutations, but also enables integration of much larger DNA sequences. We applied this advantage of the system to rapidly reprogram the antigen specificity of human T cells, which is critical for many cellular immunotherapy applications. Recent work demonstrates that chimeric antigen receptors (CARs) have enhanced efficacy when they are genetically encoded in the endogenous *TCR* locus using CRISPR/Cas9 gene cutting and an adeno-associated virus vector as a repair template^8^. Targeting of specific TCR sequences to this locus has not been achieved and is challenging because a cell must express paired TCR alpha (TCR-α) and beta chains (TCR-β) to make a functional receptor.

We developed a strategy to replace the endogenous TCR using non-viral genome targeting to integrate an approximately 1.5 kb DNA cassette into the first exon of the TCR-α constant region *(TRAC)* (**Fig. 4a**). This cassette encoded the full-length sequence of a TCR-β separated by a self-excising 2A peptide from the variable region of a new TCR-α, which encodes the full TCR-α sequence when integrated at the endogenous *TRAC* exon (**Extended Data Fig. 18**). To test this strategy, we introduced a TCR-β and TCR-α pair (1G4) recognizing the NY-ESO-1 tumour antigen into the *TRAC* locus of polyclonal T cells isolated from healthy human donors^33^. Antibody staining for total TCR-α/β expression and NY-ESO-1-MHC dextramer staining for the NY-ESO-1 TCR expression revealed that non-viral genome targeting enabled reproducible replacement of the endogenous TCR in both CD8 and CD4 primary human T cells (**Fig. 4b** and **Extended Data Fig. 19**). The majority of T cells that did not show NY-ESO-1 TCR expression were TCR knockouts (**Fig. 4b**), presumably due to NHEJ events induced by the Cas9-mediated double-stranded breaks in *TRAC* exon 1. Up to ~70% of resulting TCR-positive cells specifically recognized the NY-ESO-1 dextramer.

**Figure 4:**
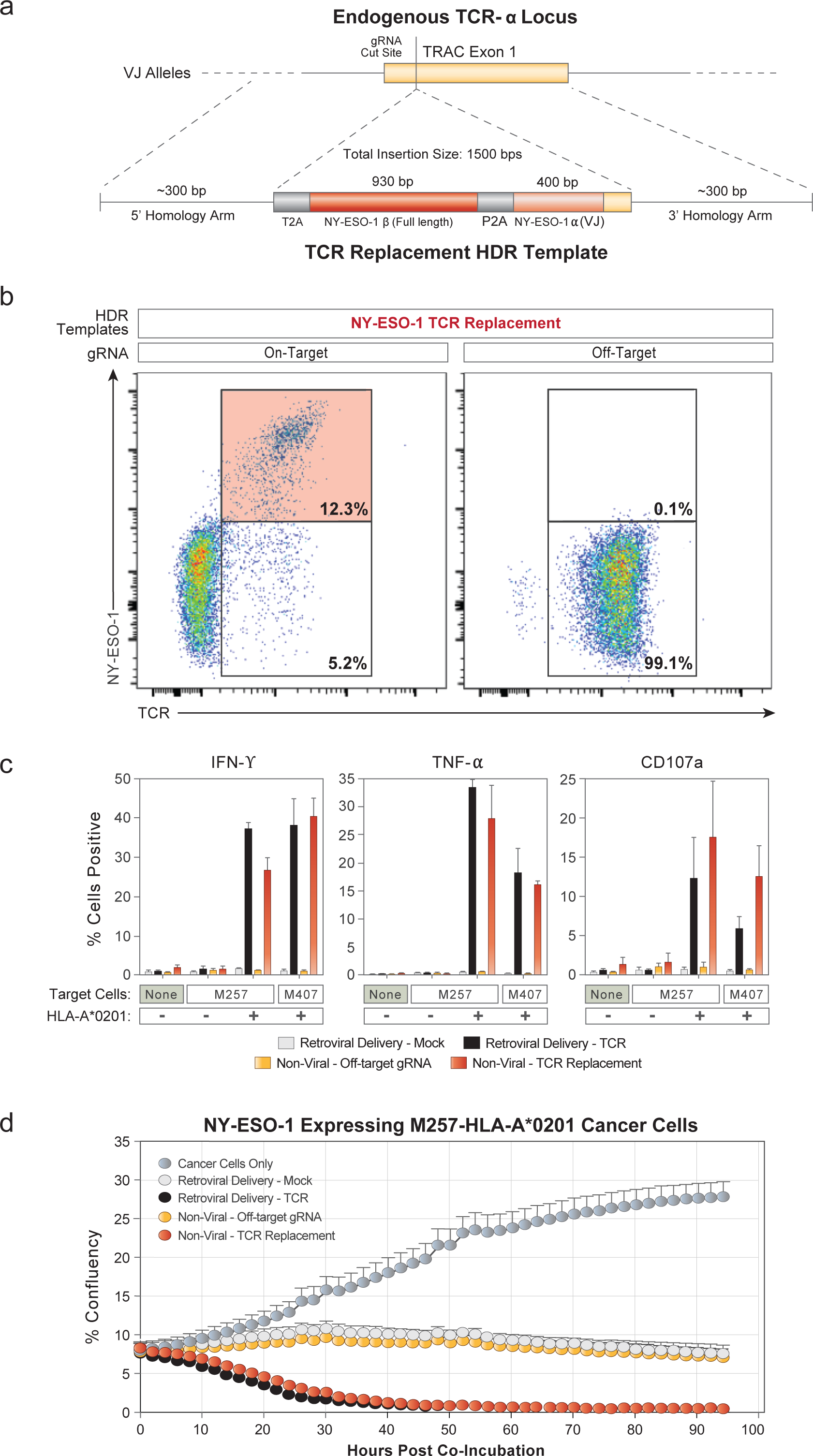
Replacement of the endogenous TCR by non-viral genome targeting. **a,** A non-viral dsDNA HDR template was used to replace the endogenous T cell receptor to reprogram antigen specificity. A sequence encoding a full-length TCR-ß receptor and the variable regions of a TCR-a receptor with a defined specificity was integrated into the first exon of the TCR constant region (TRAC), separated by self-excision peptides. A new TCR with user-defined specificity (in this case the 1G4 TCR specific for the NY-ESO-1 cancer testis antigen) is then expressed under endogenous regulatory control. **b,** Non-viral genome targeting successfully replaced the endogenous TCR repertoire in primary human CD8+ T cells. Staining with an HLA-A*0201 class I MHC restricted NY-ESO-1 peptide dextramer as well as a total TCRaß antibody reveals a robust population of NY-ESO-1 TCR+ T cells only with the presence of an on-target gRNA. Up to ~70% of resulting TCR-positive cells recognized the NY-ESO-1 dextramer. **c,** Functional antigen specific cytokine production and degranulation were observed in CD8+ T cells with the replaced TCR. 14 days after electroporation, IFN-L, TNF-a, and CD107a expression was assayed 6 hours after co-culture with target melanoma cell lines. Cytokine release and degranulation in NY-ESO-1 TCR+ cells was similar to positive control retrovirally transduced cells (from the same blood donor) expressing the same NY-ESO-1 TCR, and was specific to cell lines with the matched HLA-A*0201 class I MHC allele. **d,** Antigen specific killing of target cell by T cells with non-viral TCR replacement. T cells were cocultured with the indicated target melanoma cell line (5:1 ratio) 14 days after electroporation and *in vitro* target cell killing was assayed by time-lapse microscopy on an Incucyte microscope. Displayed data are averages and standard deviations from technical triplicates and are representative of two blood donors (**b, c, d**).

Next, we assessed the tumour antigen-specific function of targeted human T cells. When the targeted T cells were co-cultured with two different NY-ESO-1 + melanoma cell lines, M257 and M407, the modified T cells robustly and specifically produced IFN-L and TNF-α and induced T cell degranulation (measured by CD107a surface expression) **(Fig. 4c**). Cytokine production and degranulation only occurred when the NY-ESO-1 TCR transgenic T cells were exposed to cell lines expressing the appropriate HLA-A*0201 class I MHC allele required to present the cognate NY-ESO-1 peptide. Both the CD8 and CD4 T cell response was consistent across healthy donors, and was comparable to the response of T cells from the same healthy donor in which the NY-ESO-1 TCR was transduced by gamma retrovirus and heterologously expressed using a viral promoter (**Fig. 4c** and **Extended Data Fig. 19**).

Finally, we confirmed the anti-tumour cell killing ability of T cells following nonviral endogenous TCR replacement. NY-ESO-1 TCR knock-in T cells rapidly killed target M257-HLA-A*0201 cancer cells at similar rates to the positive control, retrovirally transduced T cells (**Fig. 4d**). Killing was NY-ESO-1 antigen-specific, consistent across donors, and dependent on the T cells being modified using both the correct gRNA and HDR template (**Extended Data Fig. 20**).

Our therapeutic gene editing in human T cells is a process that takes only a short time from target selection to production of the genetically modified T cell product. In approximately one week, novel guide RNAs and DNA repair templates can be designed, synthesized, and the DNA integrated into primary human T cells that remain viable, expandable, and functional. The whole process and all required materials can be easily adapted to good manufacturing practices (GMP) for clinical use. Avoiding the need for viral vectors will accelerate research and clinical applications, reduce the cost of genome targeting, and potentially improve safety.

Looking forward, the technology could be used to “rewire” complex molecular circuits in human T cells. Multiplexed integration of large functional sequences at endogenous loci should allow combinations of coding and non-coding elements to be corrected, inserted, modified, and rearranged. Much work remains to be done to improve our understanding endogenous T cell circuitry if we hope to build synthetic circuits.

Rapid and efficient non-viral tagging of endogenous genes in primary human cells will facilitate live-cell imaging and proteomic studies to decode T cell programs. Non-viral genome targeting provides an approach to re-write these programs in cells for the next generation of immunotherapies.

## METHODS

### Data reporting

No statistical methods were used to predetermine sample size. The experiments were not randomized and the investigators were not blinded to allocation during experiments and outcome assessment.

### Antibodies

All antibodies used in the study for fluorescence activated cell sorting, flow cytometry, and cellular stimulations are listed in **Supplementary Table 1**.

### Guide RNAs

All guide RNAs used in the study are listed in **Supplementary Table 2.**

### Isolation of human primary T cells for gene targeting

Primary human T cells were isolated from healthy human donors either from fresh whole blood samples, residuals from leukoreduction chambers after Trima Apheresis (Blood Centers of the Pacific), or leukapheresis products (StemCell). Peripheral blood mononuclear cells (PBMCs) were isolated from whole blood samples by Ficoll centrifugation using SepMate tubes (STEMCELL, per manufacturer’s instructions). T cells were isolated from PBMCs from all cell sources by magnetic negative selection using an EasySep Human T Cell Isolation Kit (STEMCELL, per manufacturer’s instructions). Unless otherwise noted, isolated T cells were stimulated as described below and used directly (fresh). When frozen cells were used, previously isolated T cells that had been frozen in Bambanker freezing medium (Bulldog Bio) per manufacturer’s instructions were thawed, cultured in media without stimulation for 1 day, and then stimulated and handled as described for freshly isolated samples. Fresh blood was taken from healthy human donors with their informed consented under a protocol approved by the UCSF Committee on Human Research (CHR #13-11950). Patient samples used for gene editing were obtained under a protocol approved by the Yale Internal Review Board (IRB). PBMCs for retroviral transduction experiments and corresponding TCR knockins were obtained from leukapheresis products of healthy donors. The leukapheresis product were collected either under UCLA Institutional Review Board (IRB) approval #10-001598 or purchased from AllCells, LLC.

### Primary human T cell culture

Unless otherwise noted, bulk T cells were cultured in XVivo15 medium (STEMCELL) with 5% Fetal Bovine Serum, 50 mM 2-mercaptoethanol, and 10 mM N-Acetyl L-Cystine. Immediately following isolation, T cells were stimulated for 2 days with anti-human CD3/CD28 magnetic dynabeads (ThermoFisher) at a beads to cells concentration of 1:1, along with a cytokine cocktail of IL-2 at 200 U/mL (UCSF Pharmacy), IL-7 at 5 ng/mL (ThermoFisher), and IL-15 at 5 ng/mL (Life Tech). Following electroporation, T cells were cultured in media with IL-2 at 500 U/mL. Throughout the culture period T cells were maintained at an approximate density of 1 million cells per mL of media. Every 2-3 days post-electroporation additional media was added, along with additional fresh IL-2 to bring the final concentration to 500 U/mL, and cells were transferred to larger culture vessels as necessary to maintain a density of 1 million cells/mL.

### RNP production

RNPs were produced by complexing a two-component gRNA to Cas9, as previously described^14^. Briefly, crRNAs and tracrRNAs were chemically synthesized (Dharmacon, IDT), and recombinant Cas9-NLS, D10A-NLS, or dCas9-NLS were recombinantly produced and purified (QB3 Macrolab). Lyophilized RNA was resuspended in 10 mM Tris-HCL (7.4 pH) with 150 mM KCl at a concentration of 160 μM, and stored in aliquots at -80C. crRNA and tracrRNA aliquots were thawed, mixed 1:1 by volume, and annealed by incubation at 37C for 30 min to form an 80 μNM gRNA solution. Recombinant Cas9 or the D10A Cas9 variant were stored at 40 μM in 20 mM HEPES-KOH pH 7.5, 150 mM KCl, 10% glycerol, 1 mM DTT, were then mixed 1:1 by volume with the 80 μM gRNA (2:1 gRNA to Cas9 molar ratio) at 37C for 15 min to form an RNP at 20 μM. RNPs were electroporated immediately after complexing.

### Double stranded DNA HDRT production

Novel HDR sequences were constructed using Gibson Assemblies to insert the HDR template sequence, consisting of the homology arms (commonly synthesized as gBlocks from IDT) and the desired insert (such as GFP) into a cloning vector for sequence confirmation and future propagation. These plasmids were used as templates for highoutput PCR amplification (Kapa Hotstart polymerase). PCR amplicons (the dsDNA HDRT) were SPRI purified (1.0X) and eluted into a final volume of 3 μL H2O per 100 μL of PCR reaction input. Concentrations of HDRTs were determined by nanodrop using a 1:20 dilution. The size of the amplified HDRT was confirmed by gel electrophoresis in a 1.0% agarose gel. All homology directed repair template sequences used in the study, both dsDNA and ssDNA, are listed in **Supplementary Table 2.**

### Single stranded DNA HDRT production by exonuclease digestion

To produce long ssDNA as HDR templates, the DNA of interest was amplified via PCR using one regular, non-modified PCR primer and a second phosphorylated PCR primer. The DNA strand that will be amplified using the phosphorylated primer, will be the strand that will be degraded using this method. This makes it possible to prepare either a single-stranded sense or single-stranded antisense DNA using the respective phosphorylated PCR primer. To produce the ssDNA strand of interest, the phosphorylated strand of the PCR product was degraded by treatment with two enzymes, Strandase Mix A and Strandase Mix B, for 5 minutes (per 1kb) at 37C, respectively. Enzymes were deactivated by a 5 minute incubation at 80C. The resulting ssDNA HDR templates were SPRI purified (1.0X) and eluted in H2O. A more detailed protocol for the Guide-it™ Long ssDNA Production System (Takara Bio USA, Inc. #632644) can be found at the manufacturer’s website.

### Single stranded DNA HDRT production by reverse synthesis

ssDNA HDR templates were synthesized by reverse transcription of an RNA intermediate followed by hydrolysis of the RNA strand in the resulting RNA:DNA hybrid product, as described^25^. Briefly, the desired HDR donor was first cloned downstream of a T7 promoter and the T7-HDR donor sequence amplified by PCR. RNA was synthesized by *in vitro* transcription using HiScribe T7 RNA polymerase (New England Biolabs) and reverse-transcribed using TGIRT-III (InGex). Following reverse transcription, NaOH and EDTA were added to 0.2 M and 0.1 M respectively and RNA hydrolysis carried out at 95C for 10 min. The reaction was quenched with HCl, the final ssDNA product purified using Ampure XP magnetic beads (Beckman Coulter) and eluted in sterile RNAse-free H2O. ssDNA quality was analysed by capillary electrophoresis (Bioanalyzer, Agilent).

### Primary T cell electroporation

RNPs and HDR templates were electroporated 2 days following initial T cell stimulation. T cells were harvested from their culture vessels and magnetic anti-CD3/anti-CD28 dynabeads were removed by placing cells on an EasySep cell separation magnet for 2 minutes. Immediately prior to electroporation, de-beaded cells were centrifuged for 10 minutes at 90g, aspirated, and resuspended in the Lonza electroporation buffer P3 using 20 μL buffer per one million cells. For optimal editing one million T cells were electroporated per well using a Lonza 4D 96-well electroporatior system with pulse code EH115. Alternate cell concentrations from 200,000 up to 2 million cells per well resulted in lower transformation efficiencies. Alternate electroporation buffers were used as indicated, but had different optimal pulse settings (EO155 for OMEM buffer). Unless otherwise indicated, 2.5 μL of RNPs (50 pmols total] were electroporated, along with 2 μL of HDR Template at 2 μg/μL (4 μg HDR Template total).

The order of cell, RNP, and HDRT addition appeared to matter **(Extended Data Fig, 1)**. For 96-well experiments, HDRTs were first aliquoted into wells of a 96-well polypropylene V-bottom plate. RNPs were then added to the HDRTs and allowed tc incubate together at RT for at least 30 seconds. Finally, cells resuspended in electroporation buffer were added, briefly mixed by pipetting with the HDRT and RNP, and 24 μLs of total volume (cells + RNP + HDRT) was transferred into a 96 well electroporation cuvette plate. Immediately following electroporation, 80 μLs of prewarmed media (without cytokines) was added to each well, and cells were allowed to rest for 15 minutes at 37C in a cell culture incubator while remaining in the electroporation cuvettes. After 15 minutes, cells were moved to final culture vessels.

### Flow cytometry and cell sorting

Flow cytometric analysis was performed on an Attune NxT Accustic Focusing Cytometei (ThermoFisher) or an LSRII FACs machine (BD). Fluorescence activated cell sorting was performed on an Aria II Cell Sorter (BD). Surface staining for flow cytometry anc cell sorting was performed by pelleting cells and resuspending in 25 μL of FACS Buffer (2% FBS in PBS) with antibodies at the indicated concentrations (**Supplementary Table 1**) for 20 minutes at 4C in the dark. Cells were washed once in FACS buffer before resuspension.

### Confocal microscopy

Samples were prepared by drop casting 10 μl of a solution of suspended live T cells onto a 3×1” microscope slide onto which a 25 mm^2^ coverslip was placed. Imaging was performed on an upright configuration Nikon A1r laser scanning confocal microscope. Excitation was achieved through a 488 nm OBIS laser (Coherent). A long working distance (LWD) 60x Plan Apo 1.20 NA water immersion objective was used with additional digital zoom achieved through the NIS-Elements software. Images were acquired under “Galvano” mirror settings with 2x line averaging enabled and exported as TIFF to be analyzed in FIJI (ImageJ, NIH).

### CUT&RUN

CUT&RUN was performed using epitope-tagged primary human T cells 11 days after electroporation and 4 days after re-stimulation with anti-CD3/anti-CD28 dynabeads (untagged cells were not electroporated). Approximately 20% and 10% of electroporated cells showed GFP-BATF expression as determined by flow cytometry in donor 1 and donor 2 samples, respectively. CUT&RUN was performed as described^16^, using anti-GFP (ab290), anti-BATF (sc-100974), and rabbit anti-mouse (ab46540) antibodies. Briefly, 6 million cells (30 million cells for anti-GFP CUT&RUN in GFP-BATF-containing cells) were collected and washed. Nuclei were isolated and incubated rotating with primary antibody (GFP or BATF) for 2 hours at 4C. BATF CUT&RUN samples were incubated an additional hour with rabbit anti-mouse antibody. Next, nuclei were incubated with proteinA-micrococcal nuclease (kindly provided by the Henikoff lab) for one hour at 4C. Nuclei were equilibrated to 0C and MNase digestion was allowed to proceed for 30 minutes. Solubilized chromatin CUT&RUN fragments were isolated and purified. Paired-end sequencing libraries were prepared and analysed on Illumina Nextseq machines and sequencing data was processed as described^16^. For peak calling and heatmap generation, reads mapping to centromeres were filtered out.

### TLA sequencing and analysis

TLA sequencing was performed by Cergentis as previously described^20^. Similarly, data analysis of integration sites and transgene fusions was performed by Cergentis as previously described^20^. TLA sequencing was performed in two healthy donors, each edited at the *RAB11A* locus with either a dsDNA or ssDNA HDR template to integrate a GFP fusion (**Fig. 1b**). Sequencing reads showing evidence of primer dimers or primer bias (i.e. greater than 99% of observed reads came from single primer set) were removed.

### *In vitro* Treg suppression assay

CD4+ T cells were enriched using the EasySep Human CD4+ T cell enrichment kit (STEMCELL Technologies). CD3+CD4+CD127loCD45RO+TIGIT+ enriched Treg-like cells from IL2RA-deficient subjects and HD as well as CD3+CD4+IL2RAhiCD127lo Tregs from IL2RA+/−individuals were sorted by flow cytometry. CD3+CD4+IL2RA-CD127+ responder T cells (Tresps) were labeled with CellTrace CFSE (Invitrogen) at 5 μM. Tregs and HD Tresps were co-cultured at a 1:1 ratio in the presence of beads loaded with anti-CD2, anti-CD3 and anti-CD28 (Treg Suppression Inspector; Miltenyi Biotec) at a 1 bead: 1 cell ratio. On days 3.5 to 4.5, co-cultures were analyzed by FACS for CFSE dilution. % inhibition is calculated using the following formula: 1 -(% proliferation with Tregs / % proliferation of stimulated Tresps without Tregs).

### Generation of retrovirally transduced PBMCs control cells

PBMCs from healthy donors were thawed, activated with CD3/CD28 beads (Invitrogen, Carlsbad, CA) at a 1:1 ratio and expanded in T cell media (RPMI supplemented with 10% fetal bovine serum, penicillin-streptomycin and 300IU/mL of hIL2). Forty-eight hours after activation, PBMCs were transduced with clinical grade MSGV-1-1G4 (NY-ESO-1 TCR transgene) retroviral vector (IUVPC, Indianapolis, IN) by spinoculation in retronectin (Clontech, Mountain View, CA) coated plates. Control mock-transduced PBMCs were generated. PBMCs were expanded for 8 days after transduction, aliquoted and stored in liquid nitrogen.

### Antigen specific TCR expression analysis

Modified PBMCs were thawed and expanded in T cell media supplemented with 300IU/mL hIL2. To assess the expression of the NY-ESO-1 TCR cells were stained with NY-ESO-1 specific (SLLMWITQC) dextramer-PE (Immundex, Copenhagen, Denmark), Negative dextramer (Immudex, Copenhagen, Denmark) was used as a negative control.

### T cell activation and cytokine production analysis

Melanoma cell lines were established from the biopsies of melanoma patients under the UCLA IRB approval #11-003254. Cell lines were periodically authenticated using GenePrint^®^ 10 System (Promega, Madison, WI), and were matched with the earliest passage cell lines. M257 (NY-ESO-1+ HLA-A*0201−), M257-A2 (NY-ESO-1+ HLA-A*0201+) and M407 (NY-ESO-1+ HLA-A*0201+) were cocultured 1:1 with the modified PBMCs in cytokine free media. The recommended amount per test of CD107a-APC-H7 (**Supplementary Table 1**) antibody was added to the coculture. After 1 hour, half the recommended amount of BD Golgi Plug and BD Golgi Stop (BD bioscience, San Jose, CA) was added to the coculture. After 6 hours, surface staining was performed followed by cell permeabilization using BD cytofix/cytoperm (BD bioscience, San Jose, CA) and intracellular staining according to manufacturer instructions (**Supplementary Table 1**). Negative dextramer and Fluorescence minus one (FMOs) staining were used as controls.

### T Cell killing assay

M257-nRFP, M257-A2-nRFP, A375-nRFP, and M407-nRFP melanoma cell lines stably transduced to express nuclear RFP (Zaretsky 2016 NEJM) were seeded approximately 16 hours before starting the coculture. Modified T cells were added at the indicated E:T ratios. All experiments were performed in cytokine free media. Cell proliferation and cell death was measured by nRFP real time imaging using an IncuCyte ZOOM (Essen, Ann Arbor, MI) for 5 days. Biological quadruplicates per condition were assessed.

## Data availability

CUT&RUN data will be publically deposited prior to publication. TLA sequencing data is available upon request.

## ACKNOWLEDGEMENTS

We thank members of the Marson lab, Chris Jeans and the QB3 MacroLab, Kyle Marchuk and the UCSF Biological Imaging Development Center, Jeffrey Bluestone and Jacob Corn for suggestions and technical assistance. We thank Lonza for technical assistance and providing reagents to test electroporation conditions. This research was supported by NIH grants DP3DK111914-01 (A.M.), P50GM082250 (A.M.) and R35 CA197633 (A.R.), a grant from the Keck Foundation (A.M.), gifts from Jake Aronov and Galen Hoskin (A.M.), a gift from the Jeffrey Modell Foundation, and a National Multiple Sclerosis Society grant (A.M.; CA 1074-A-21). T.L.R. and J.H. were supported by the UCSF Medical Scientist Training Program (T32GM007618). T.L.R. was supported by the UCSF Endocrinology Training Grant (T32 DK007418). A.M. holds a Career Award for Medical Scientists from the Burroughs Wellcome Fund and is an investigator at the Chan Zuckerberg Biohub. C.P.S., J.S. and A.R. are funded by the Ressler Family Fund. A.R. is a member of the Parker Institute for Cancer Immunotherapy. The UCSF Flow Cytometry Core was supported by the Diabetes Research Center grant NIH P30 DK063720. SHH and AMF were supported by the NIH Intramural Program, Center for Cancer Research, National Cancer Institute.

## AUTHOR INFORMATION

### Contributions

T.L.R. and A.M. designed the study and wrote the manuscript. T.L.R. designed and performed all electroporation experiments. T.L.R., R.Y., E.S. J.H. J.L. V.T., D.N., and K.S. contributed to functional assays of edited T cells. R.Y. performed and analyzed CUT&RUN experiments. H.L., J.W., and M.L. developed the IVT-RT ssDNA production method. H.M., M.M, Y.M, B.S, and M.H. developed the exonuclease based ssDNA production method. R.Q. and C.G. advised on the use of ssDNA. A.M.F. and S.H.H. advised on methods of DNA introduction into T cells. A.P.M. advised on integration site analysis. J.C., J.N.S., L.P., D.C, G.A.A., D.D.G., G.K., S.G., R.B., E.M., M.G.R., N.R., and K.C.H. contributed to the clinical workup of the *IL2RA* deficient family and functional assays on unedited patient T cells. T.L.R., C.P.S., E.S., A.R., and A.M. designed the endogenous TCR knock-in strategy. T.L.R., C.P.S., J.C., J.S., A.A., and A.R. performed or supervised functional assays of T cells with endogenous TCR knock-ins.

### Corresponding Author

Correspondence and requests for materials should be addressed to.

## EXTENDED DATA FIGURES AND TABLES

**Extended Data Figure 1:**
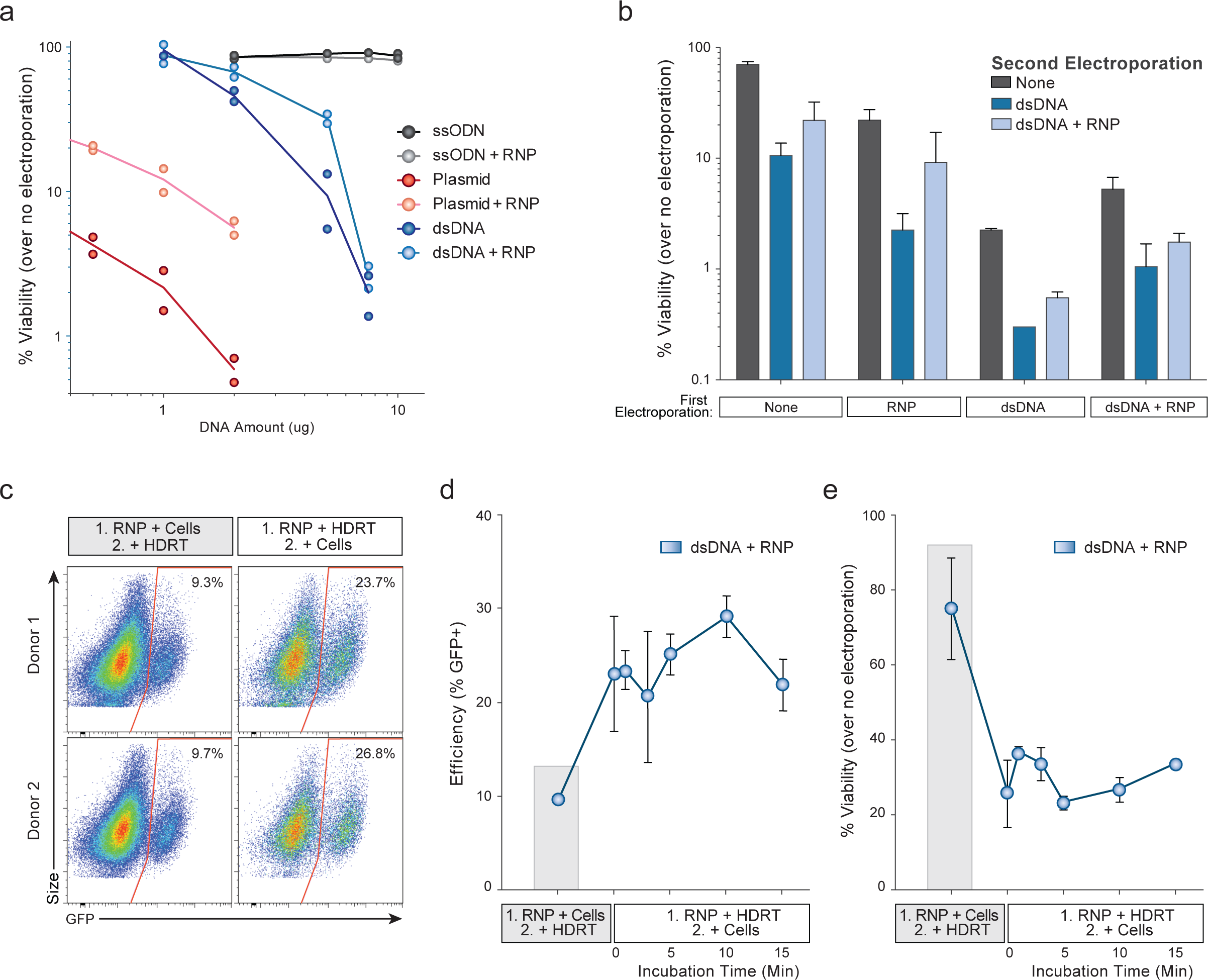
Lower reduction in cell viability when dsDNA is coelectroporated with CRISPR-Cas9 RNPs. **a,** Double-stranded (ds)DNA, both circular (plasmid) and linear, when electroporated into primary human T cells cause marked viability loss with increasing amounts of template. Co-delivery of an RNP caused less reduction in viability post electroporation. Of note, in these experiments no loss in viability was seen with short single-stranded (ss)DNA oligo donor nucleotides (ssODNs). **b,** RNPs must be delivered concurrently with DNA to see increased viability. T cells from two donors were each electroporated twice with an eight-hour rest in between electroporations. While two electroporations so closely interspersed caused a high degree of cell death, delivery of the RNP and linear dsDNA template could be delivered separately. Initial RNP electroporation did not protect from the loss of viability if dsDNA was delivered in the second round of electroporation. **c-d,** We assayed whether the order of adding reagents influenced targeting efficiency and viability. Indeed, in wells where the RNP and the DNA HDR template were mixed together prior to adding the cells (1. RNP + HDRT; 2. + Cells), no matter how long the RNP and DNA template were preincubated, there was a marked increase in targeting efficiency. **e,** Note, with the high concentration of dsDNA used in these experiment, viability was higher if the RNP and cells were mixed first and the DNA template was added immediately prior to electroporation (1. RNP + Cells; 2. + HDRT). Taken together, these data likely suggest that pre-incubation of the RNP and HDR template, even for a short period, increases the amount of DNA HDR template delivered into the cell, which increases efficiency but decreases viability. However, viability after RNP and dsDNA HDR template pre-incubation was still much higher than was observed with dsDNA HDR template electroporation by itself (**a**). Viability compared to no electroporation controls was measured 2 days following electroporation and GFP expression was measured at day 4. 5 μg of dsDNA HDR temple was used in (**c-e**). Graphs display average plus standard deviation in two donors.

**Extended Data Figure 2:**
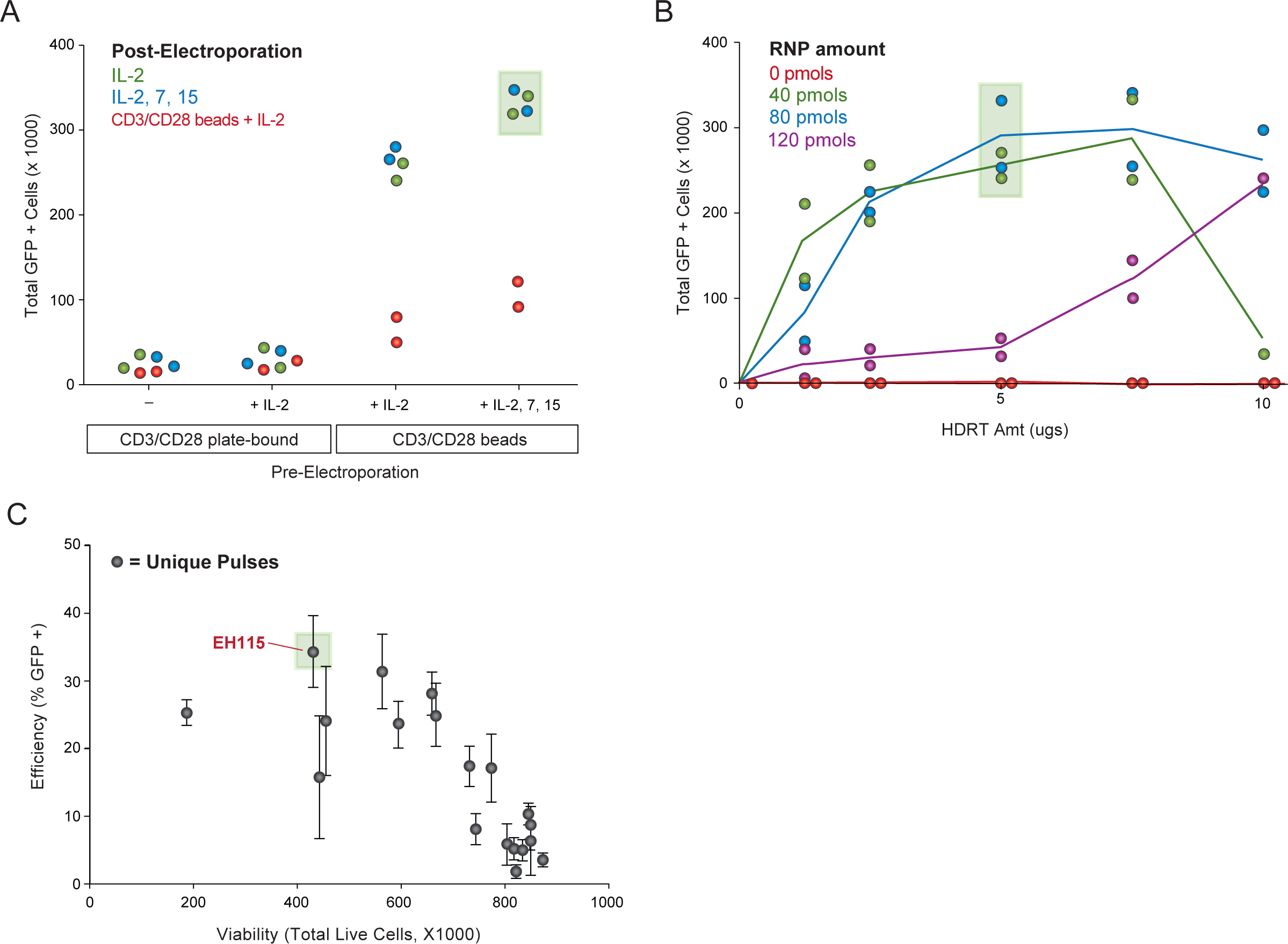
Optimization of non-viral genome targeting. **a,** Primary human T cells were cultured for 2 days using varying combinations of T cell receptor (TCR) stimulation and cytokines prior to electroporation of *RAB11A* targeting RNP and HDR template, followed by varying culture conditions post-electroporation. **b,** Among RNP and HDR template concentrations tested here, optimal GFP insertion into *RAB11A* was achieved at intermediate concentrations of the RNP and dsDNA HDRT. **c,** Arrayed testing of electroporation pulse conditions showed that, in general, conditions yielding higher HDR efficiency decreased viability. EH115 was selected to optimize efficiency, while still maintaining sufficient viability. All experiments display individual points or averages with standard deviation from two donors. Efficiency of GFP integration and viability were measured at 4-5 days post electroporation.

**Extended Data Figure 3:**
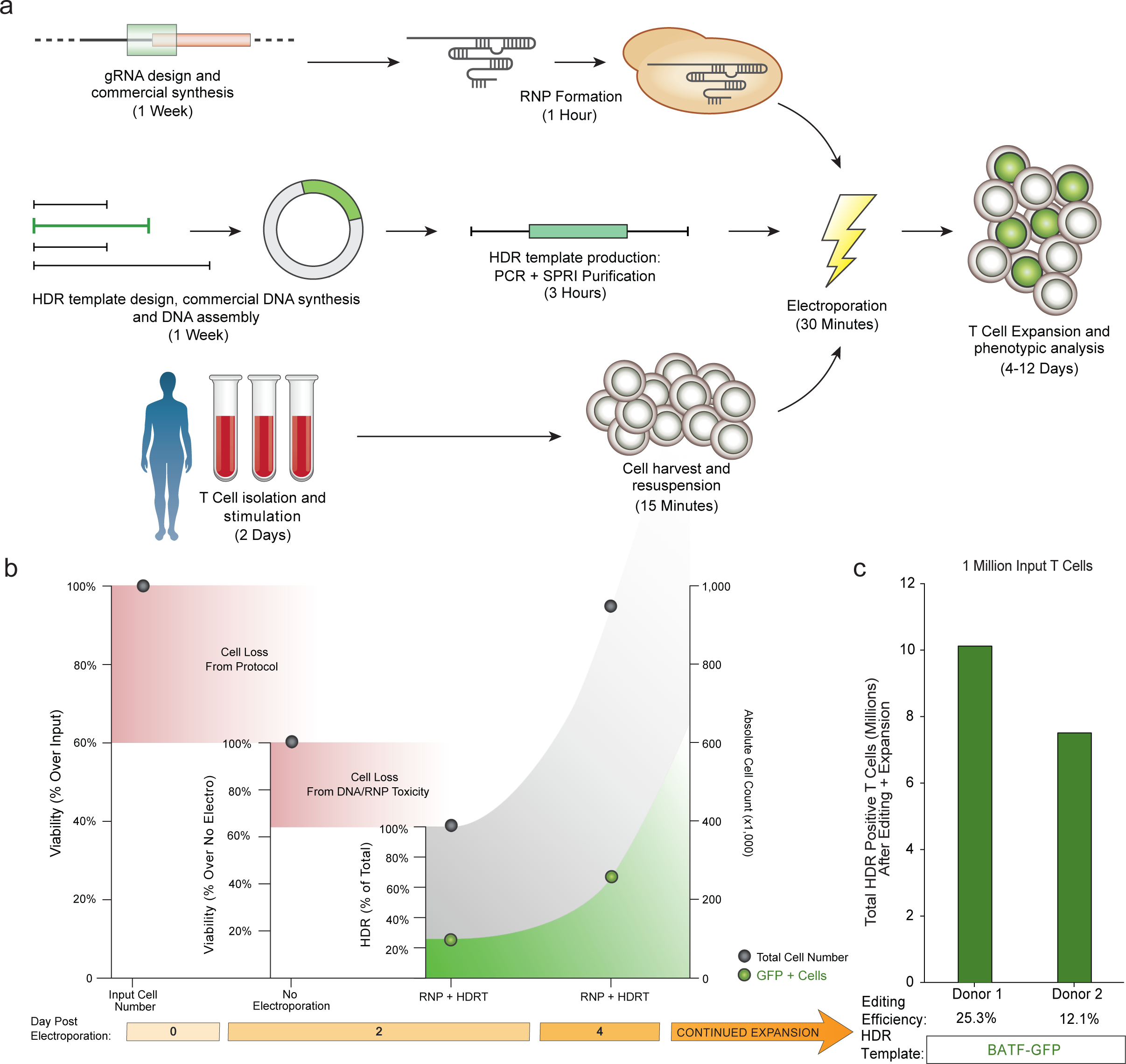
Non-viral genome targeting enables rapid and efficient genetic engineering in primary human T cells. **a,** Diagrammatic timeline of non-viral genome targeting. Approximately one week is required to design, order from commercial suppliers, and assemble any novel combination of genomic editing reagents (gRNA along with homology directed repair template). Two days prior to electroporation, primary human T cells isolated from blood or other sources (**Extended Data Fig. 4**) are stimulated. dsDNA HDR templates can be made easily by PCR followed by a SPRI purification to achieve a highly concentrated and pure product suitable for electroporation. On the day of electroporation, the gRNA (complexed with Cas9 to form an RNP), the HDR template, and harvested stimulated T cells are mixed and electroporated, a process taking approximately 1.5 hours. After electroporation, engineered T cells can be readily expanded for an additional two weeks. b, We use viability to refer to the percentage of live cells relative to an equivalent population that went through all protocol steps except for the actual electroporation (No electroporation control). We use the term efficiency to refer to the percentage of live cells in culture expressing the “knocked in” exogenous sequence (such as GFP). Finally, the total number of cells positive for the desired integration was calculated by multiplying the efficiency by the absolute cell count. Methodological changes that maximized efficiency often were not always optimal for the total number of positive cells, and vice-versa. **c,** Viability or efficiency at any single given point in a cellular therapy manufacturing pipeline is not as important as the cumulative effects on viability, efficiency, or expandability throughout the entire process. Thus the most accurate single number to assess the overall viability and efficacy for a given cell therapy production pipeline is the number of cells necessary to isolate from a patient in order to generate the ultimate therapeutic product. One million T cells isolated from two healthy donors went through the non-viral genome targeting process to integrate a GFP fusion DNA sequence at the BATF locus (**Extended Data Fig. 6**). Following expansion for an additional 12 days after electroporation (14 days after initial stimulation), 7-10 million GFP+ cells could be isolated originating from the original 1 million input cells (of course many more GFP-cells could also be isolated). This suggests that as few as 10-15 million starting T cells could be used to generate the approximately 100 million edited T cells commonly transferred in current adoptive T cell immunotherapies^34^.

**Extended Data Figure 4:**
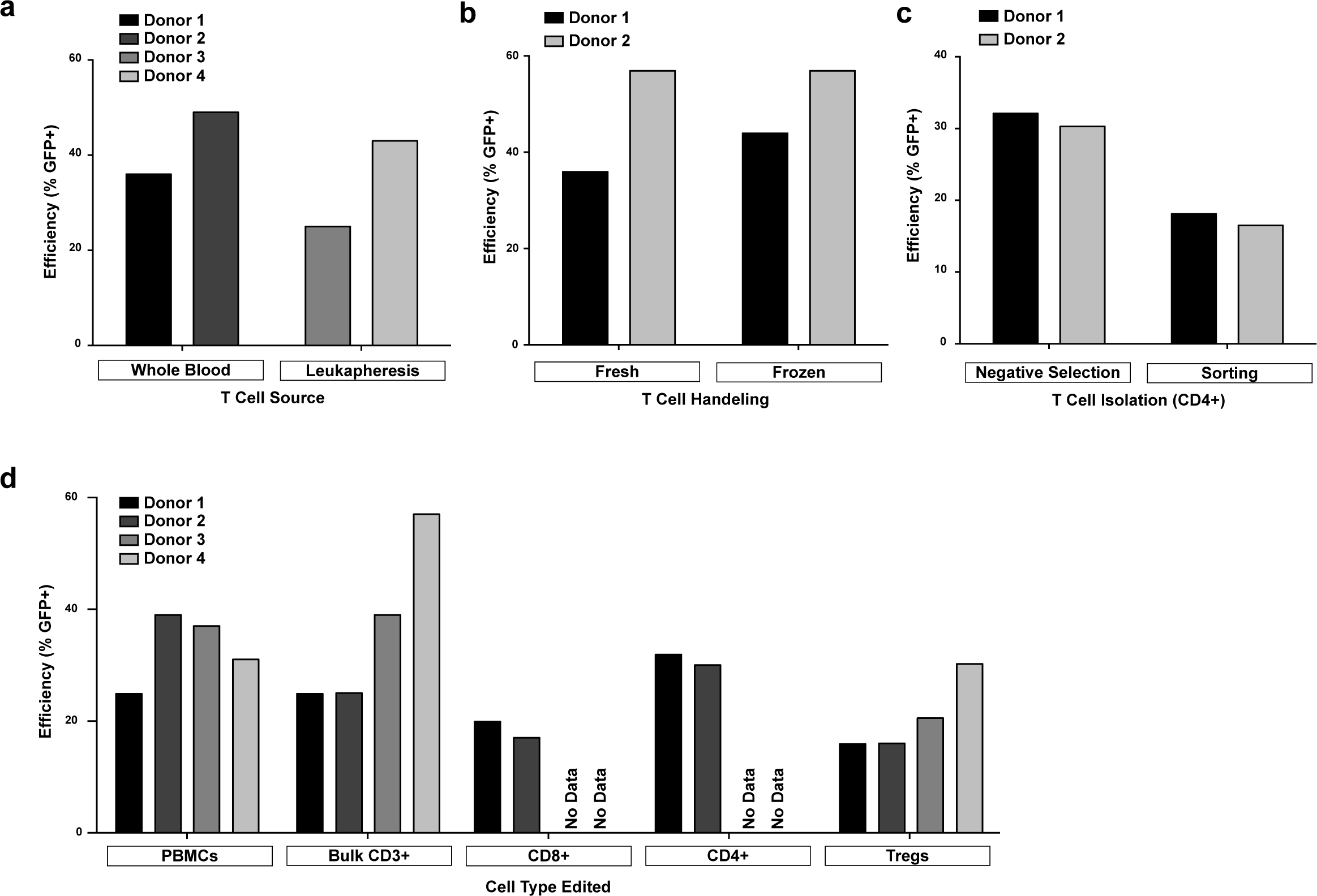
Non-viral genome targeting is consistent across T cell types, sources, handling conditions, and isolation techniques. **a,** Efficient genome targeting was accomplished with a variety of T cell processing and handling conditions used with current manufacturing protocols for cell therapies. Nonviral genome targeting of a RAB11A-GFP fusion protein using a linear dsDNA HDR template was performed in bulk CD3+ T cells isolated from either whole blood draws or by leukapheresis. **b,** Targeting was similar in bulk T cells used either freshly after isolation or cryopreserved (stored in liquid nitrogen and thawed prior to initial activation). **c,** CD4+ T cells isolated by fluorescent activated cell sorting (FACS) showed detectable GFP+ cells indicative of efficient editing, albeit at lower rates than targeting in CD4+ cells isolated by negative selection (potentially due to the added cellular stress of sorting). **d,** Using the same optimized non-viral genome targeting protocol (**Methods**), a variety of T cell types could be successfully edited, including peripheral blood mononuclear cells without any selection (T cell culture conditions cause selective growth of T cells from PBMCs). Sorted T cell subsets, CD8+, CD4+, and CD4+IL2RA+CD127lo regulatory T cells (Tregs) could be successfully targeted with GFP integration. PBMCs were cultured for two days identically to primary T cells (**Methods**). Bulk CD3+ T cells were isolated by negative enrichment. The electroporations in panel **d** only used 2 μg of dsDNA HDR template, a concentration that was later found to be less efficient than the final 4 μg (contributing to the lower efficiencies seen compared to **Fig. 1d**). RAB11A-GFP template with on-target gRNA was used in all conditions. In all displayed conditions the RAB11A-GFP template alone or with an off-target gRNA showed almost no GFP+ cells (<0.1% GFP+). Donor numbers are specific to each panel do not indicate the same donor was used across panels.

**Extended Data Figure 5:**
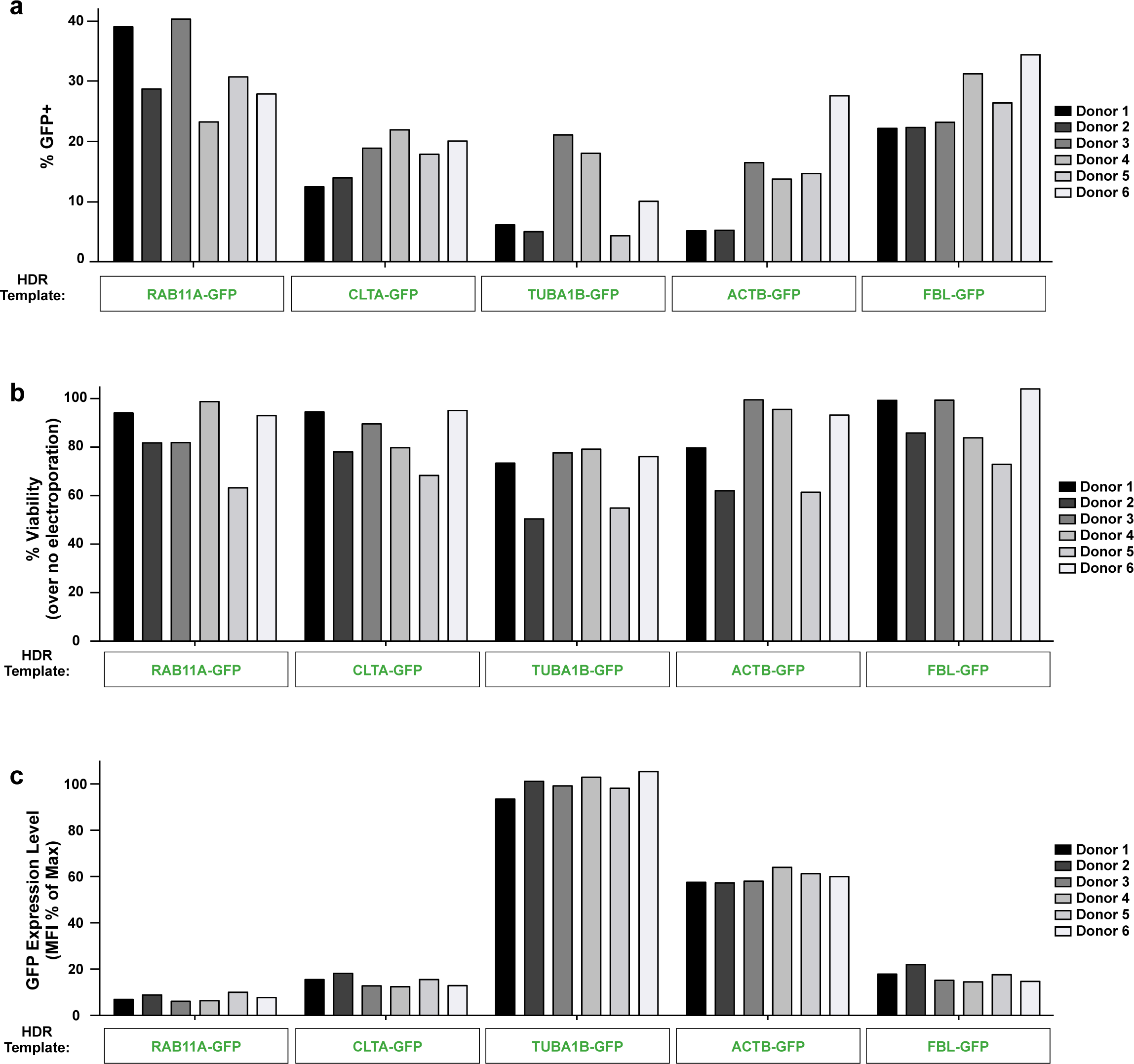
Reproducible non-viral genome targeting across different target loci. **a,** Four days after electroporation of different GFP templates along with a corresponding RNP into primary CD3+ T cells from six healthy donors, GFP expression was observed across both templates and donors. **b,** High viability post-electroporation was similarly seen across target loci. **c,** The integrated GFP fusion tags not only localize to the subcellular location of their target protein (**Fig. 2b**), but were also expressed under the endogenous gene regulation, allowing protein expression levels to be observed in living primary human T cells. Note how GFP tags of the highly expressed cytoskeletal proteins TUBA1B (beta tubulin) and ACTB (beta actin) show consistently higher levels of expression compared to the other loci targeted across six donors. GFP mean fluorescent intensity (MFI) was calculated for the GFP+ cells in each condition/donor, and normalized as a percentage of the maximum GFP MFI observed in the experiment.

**Extended Data Figure 6:**
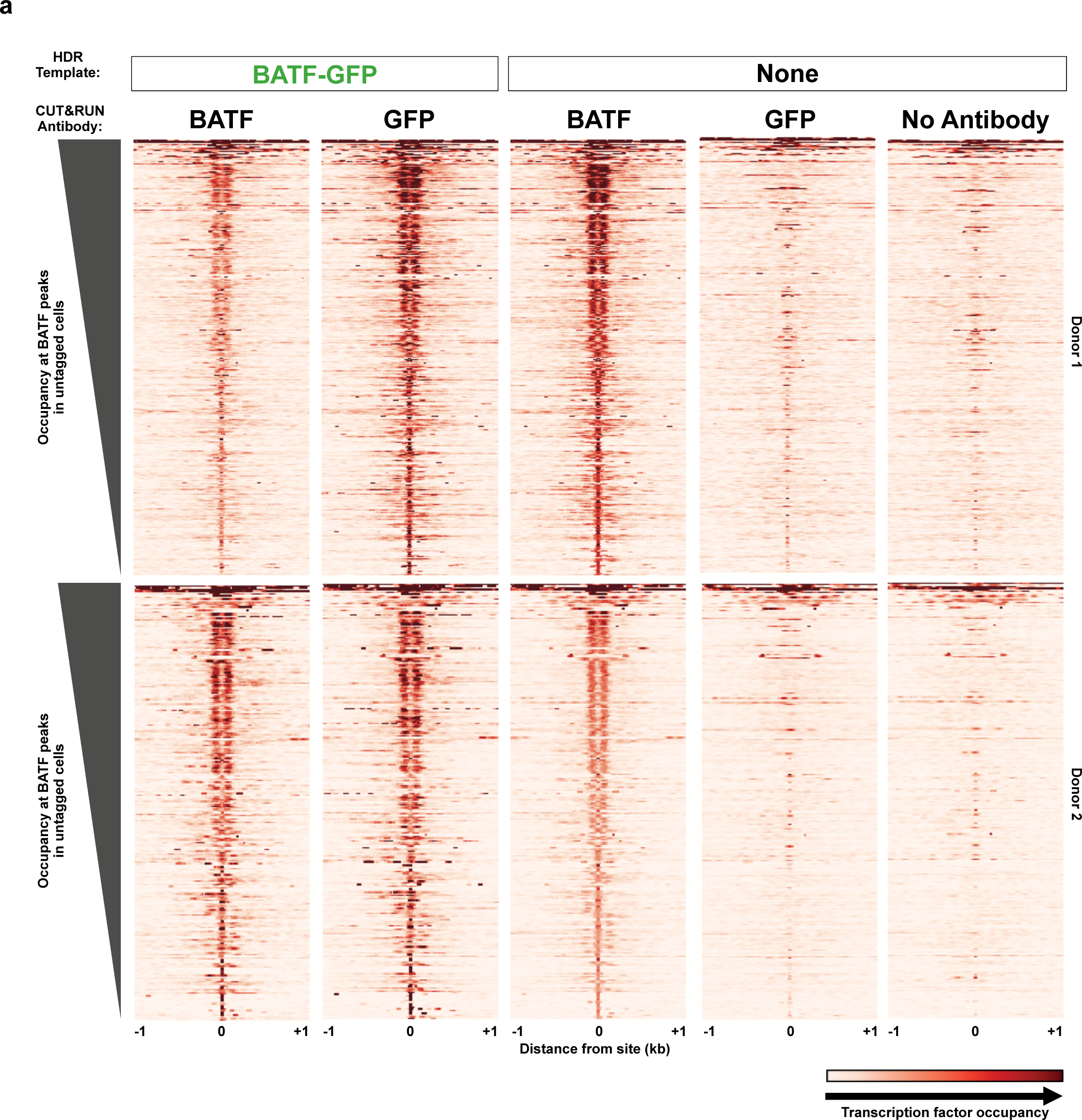
Endogenous tagging of transcription factor BATF for analysis of chromatin occupancy. **a,** Gene fusions not only permitted the imaging and analysis of expression of endogenous proteins in live cells, but also could be used for biochemical targeting of specific proteins. For example, ChIP-Seq, and more recently CUT & RUN^16^, are widely used to map transcription factor binding sites; however, these assays are often limited by the availability of effective and specific antibodies. As a proof-of-principle we used anti-GFP antibodies to perform CUT & RUN in primary T cells where the endogenous gene encoding BATF, a critical transcription factor, had been targeted to generate a GFP-fusion. Binding sites identified with anti-GFP CUT & RUN closely matched the sites identified with an anti-BATF antibody. Anti-BATF, anti-GFP, and no antibody heatmaps of CUT&RUN data obtained from primary human T cell populations electroporated with GFP-BATF fusion HDR template (untagged cells were not electroporated). Aligned CUT&RUN binding profiles for each sample were centered on BATF CUT&RUN peaks in untagged cells and ordered by BATF peak intensity in untagged cells.

**Extended Data Figure 7:**
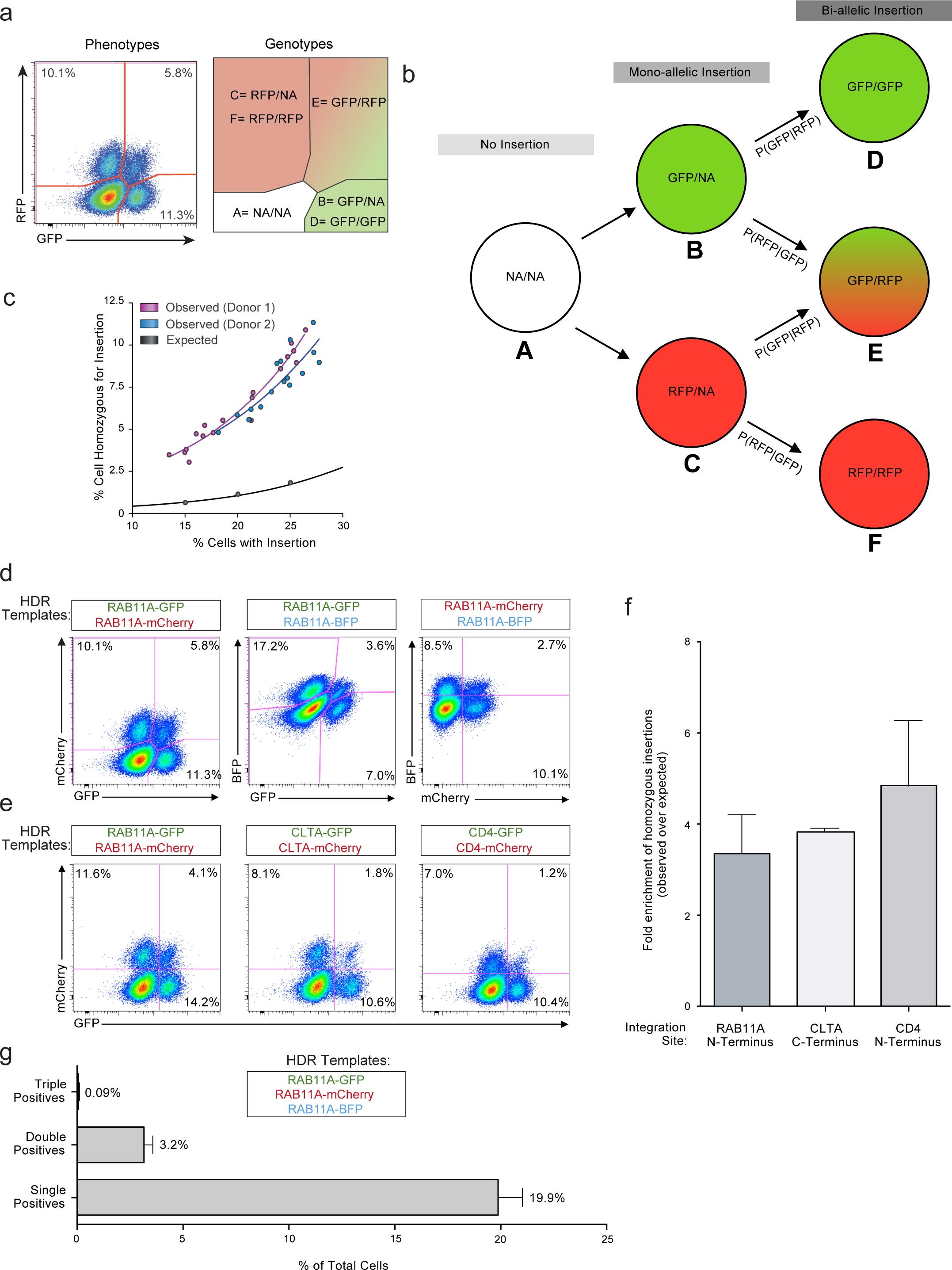
Bi-allelic HDR integrations. **a,** We wanted to confirm that we could generate cells with genome insertions in both alleles and quantify the frequency of bi-allelic modifications. Targeting the two alleles of the same gene with two distinct fluorophores would provide a way to quantify and enrich cells with bi-allelic gene modifications. The possible cellular phenotypes and genotypes when two fluorescent proteins are inserted into the same locus are displayed. Importantly, the number of cells that express both fluorescent proteins underestimates the percentage of cells with bi-allelic integrations because some cells will have inserted either GFP or mCherry on both alleles. We constructed a model to account for bi-allelic integrations of the same fluorescent protein (**Supplementary Note 1**) **b,** Diagram of bi-allelic integration model. The total percentage of cells with bi-allelic HDR integrations must be the sum of genotypes D, E, and F. While the proportion of cells with genotype E (dual fluor positives) is immediately apparent from the phenotypes, genotypes D and F are not. Our model allow for the de-convolution of the multiple genotypes in the single fluor positive phenotypes, and thus an estimation of the true percentage of cells bi-allelic for HDR. **c,** The observed level of bi-allelic integrations is higher in cells that acquired at least one integration than would be expected by chance. Individual points represent replicates where the combination of the genes encoding the fluorescent proteins was varied (either GFP + mCherry, GFP + BFP, or mCherry + BFP) as was the amount of the HDR template (3 to 6 μg). **d,** Bi-allelic HDR analysis applied across a variety of fluorophore permutations inserted into the *RAB11A* locus. **e,** Dual fluorescence bi-allelic integrations were seen across target loci. **f,** The data also suggest that cells with one mono-allelic integration are more likely to have also undergone a second targeted bi-allelic integration, and this effect was observed across three genomic loci. While the total percentage of cells with an insertion varied with the efficiency of each target site, the fold enrichment in the observed percentage of homozygous cells over that predicted by random chance was consistent across loci. **g,** Co-delivery of three fluorescent-tags targeting the *RAB11A* locus resulted in only a few cells that expressed all three fluorophores, consistent with a low rate of off-target integrations. As a max of two targeted insertions are possible (at the locus’ two alleles; assuming a diploid genome), no cells positive for all three loci should be observed (triple positives). Indeed, while large numbers of single fluorophore integrations are observed (single positives), as well as cells positive for the various permutations of two fluorophores (double positives), there is a 30 fold reduction in the number of triple positive cells compared to double positives. All flow cytometric analysis of fluorescent protein expression shown here was performed 4 days following electroporation. All displays are representative of multiple technical replicates from at least two healthy human donors. Graphs display mean + standard deviation of two donors.

**Extended Data Figure 8:**
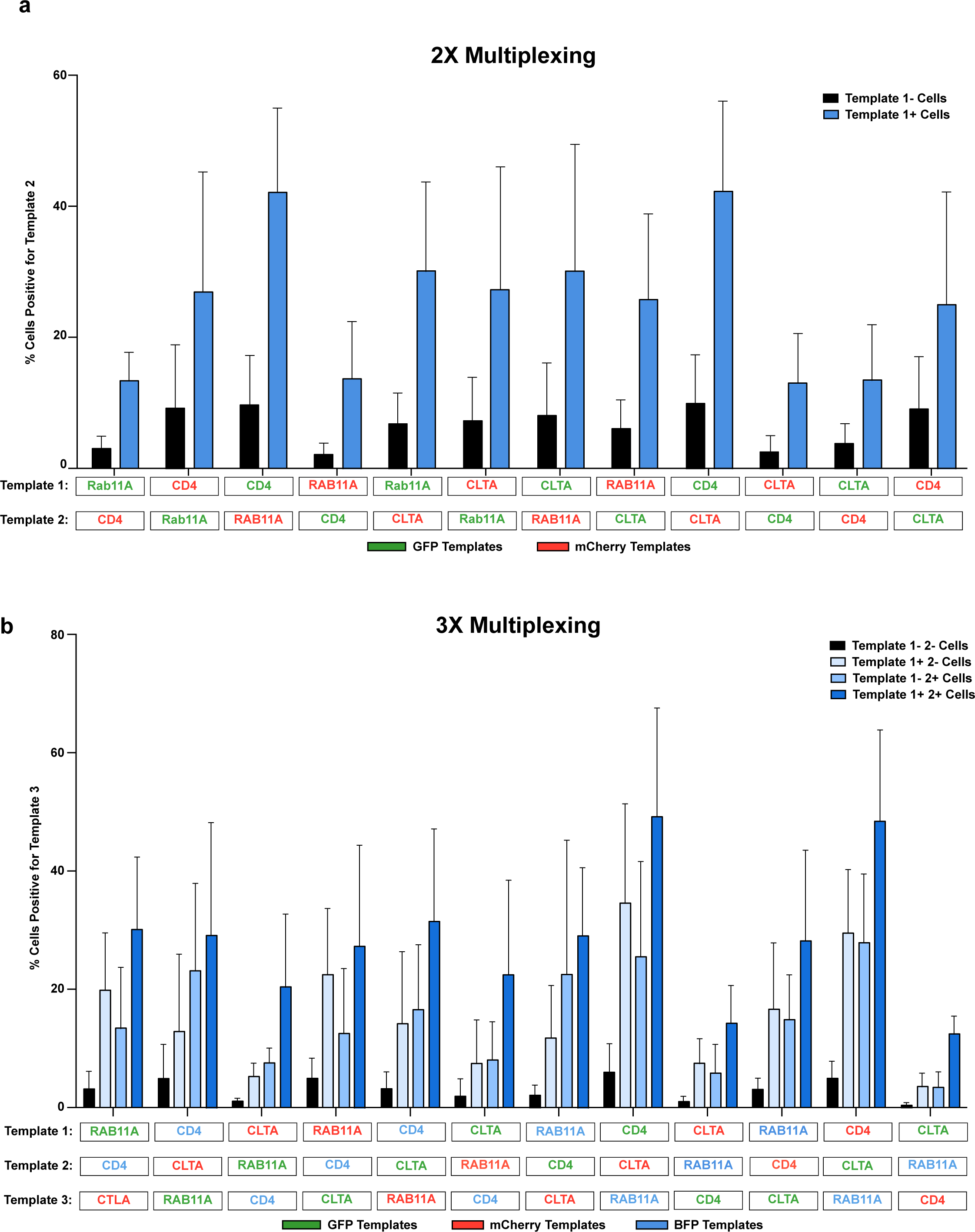
Multiplexed HDR integrations. **a,** Multiplex editing of combinatorial sets of genomic sites would support expanded research and therapeutic applications^17^. We tested whether multiple HDR templates could be co-delivered along with multiple RNPs to generate primary cells in which more than one locus was modified. Primary human T cells with two modifications were enriched by gating on the cells that had at least one modification, and this effect was consistent across multiple combinations of genomic loci. HDR template permutations from a set of six dsDNA HDR templates (targeting RAB11A, CD4, and CLTA; each site with GFP or RFP) were electroporated into CD3+ T cells isolated from healthy human donors. Four days after electroporation of the indicated two HDR templates along with their two respective on-target RNPs, the percentage of cells positive for each template was analysed when gating on cells either positive or negative for the other template. Not only was two-template multiplexing possible across a variety of template combinations, but gating on cells positive for one template (Template 1+ Cells, Blue) yielded an enriched population of cells more likely to be positive for the second template compared to cells negative for the first (Template 1-Cells, Black). 2 μg of each template, along with 30 pmols of each associated RNP, were electroporated for dual multiplexing experiments. **(B)** We were also able to achieve triple gene targeting and could significantly enrich for cells that had a third modification by gating on the cells with two targeted insertions, an effect again consistent across target genomic loci. 1.5 μg of each template (4.5 μg total) were electroporated together with 20 pmols of each corresponding RNP (60 pmols total). Displayed data are means + standard deviation from multiple technical replicates from two healthy human donors.

**Extended Data Figure 9:**
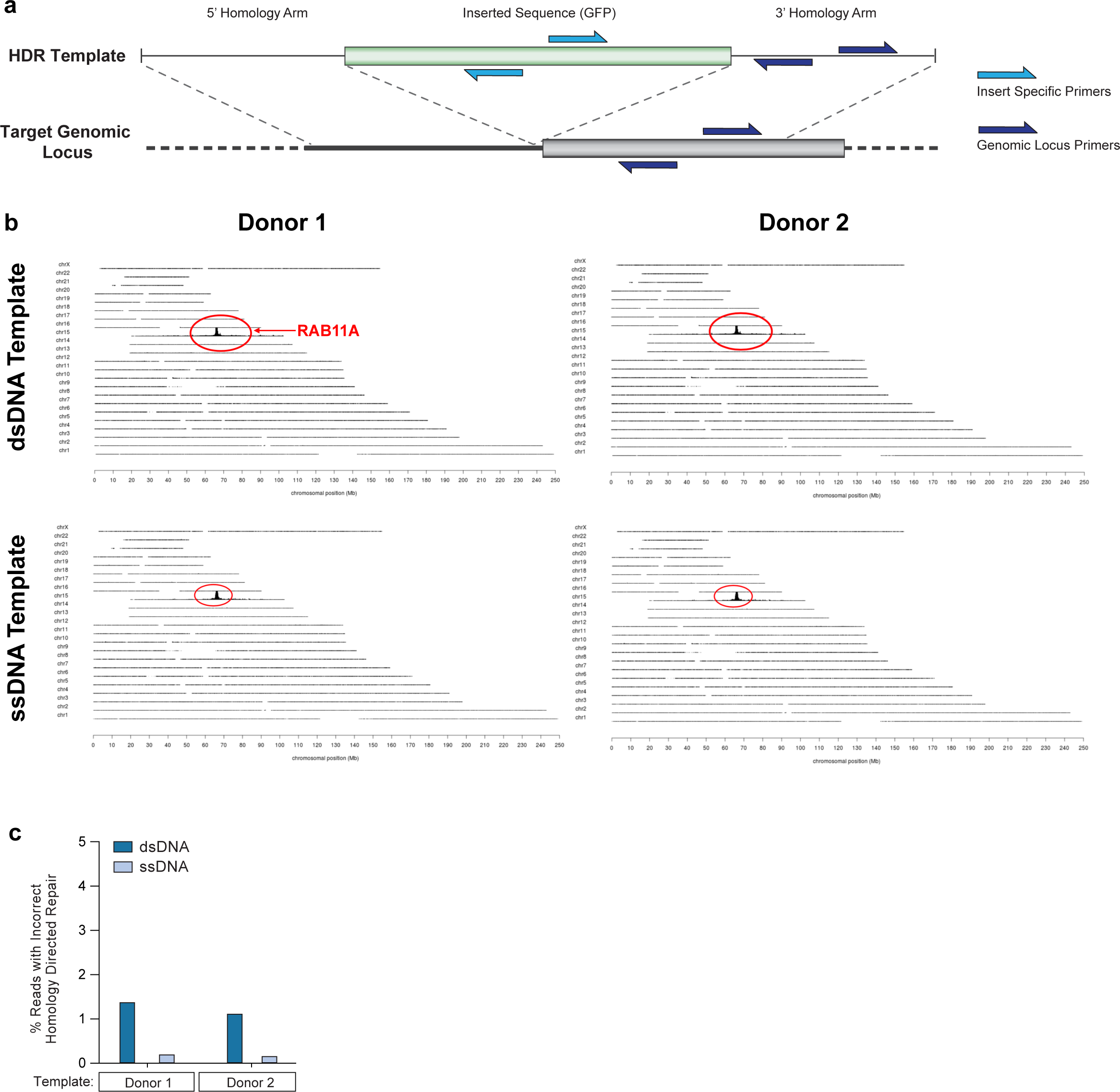
Targeted Locus Amplification sequencing reveals that off-target integrations are extremely rare with non-viral genome targeting. **a,** Diagram of sequencing primers used for Targeted Locus Amplification (TLA) off-target sequencing. Sequencing was performed with two sets of primers, one exclusive to the inserted HDR sequence (amplifying off of GFP), and one that inclusive both to the inserted HDR sequence and the genomic locus (amplifying off of the sequence contained in the 3’ Homology arm). **b,** Results of targeted locus amplification sequencing. No off-target integration sites were identified (assays limit of detection in these assays is 1% of alleles) with either a dsDNA or ssDNA HDR template in two healthy donors. The on-target *RAB11A* locus on chromosome 15 is indicated in red. **c,** Only one incorrect integration outcome at the target *RAB11A* locus was observed. The frequency of incorrect integrations at the locus could be reduced frequencies using a ssDNA HDR template in two human blood donors.

**Extended Data Figure 10:**
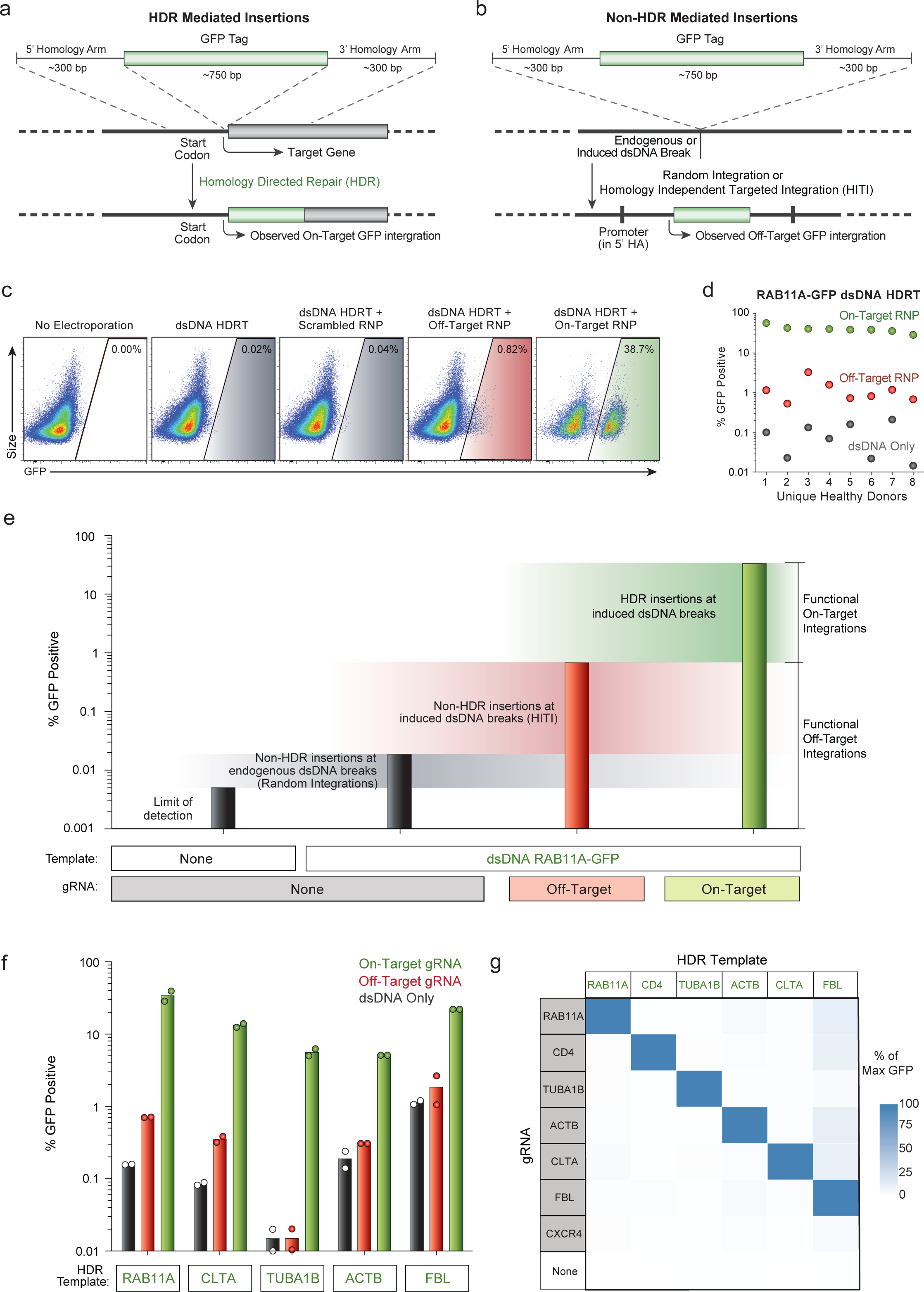
Single cell quantification of rare functional off-target integration events. **a,** Diagram of HDR mediated insertions at the N-terminus of a target locus. The homology arms specify the exact sequence where the insert (a GFP tag in this case) will be inserted, allowing for scarless integration of exogenous sequences. As a GFP fusion protein is created, GFP fluorescence will be seen as a result of this on-target integration, dependent on an RNP cutting adjacent to the integration site. **b,** dsDNA can be integrated via homology-independent repair mechanisms at off-target sites through either random integration at naturally occurring dsDNA breaks, or potentially at induced double stranded breaks, such as those at the off-target cut sites of the RNP. This effect can be harnessed to allow for targeted integration of a dsDNA sequence at a desired induced dsDNA break in quiescent cell types lacking the ability to do HDR, but crucially the entire seqeunce of the dsDNA template is integrated, including any homology arms. In the case that the homology arms contain a promoter sequence (such as for N-terminal fusion tags), these off target integrations can drive observable expression of the inserted sequence without the desired correct HDR insertion. **c,** We looked for unintended non-homologous integrations with the non-viral system using an N-terminal GFP-RAB11A fusion construct that contained the endogenous *RAB11A* promoter sequence within its 5’ homology arm. This construct can express GFP at off-target integration sites, which allowed us to assay for off-target events at the single cell level using flow cytometry. Inclusion of a gRNA designed to cut a genome region that is not the homologous region to the targeting sequence can be used to measure integration at an off-target cut site. **d,** While efficient GFP expression depended on pairing the HDR template with the correct gRNA targeting that site, rare GFP+ cells were observed when dsDNA HDR templates were delivered either alone (~0.1%) or with an "off target" Cas9 RNP (~1%). **e,** Quantification of different types of functional off-target integrations. The increase in the percentage of fluorescent cells over the limit of detection when the template alone is electroporated likely represents random integrations at naturally occurring dsDNA breaks (although cut-independent integration at the homology site is also possible in theory). Not every off-target integration will yield fluorescent protein expression, but the relative differences in functional off-target expression between different templates and editing conditions can be assayed. Inclusion of an RNP targeting *CXCR4* dramatically increases the observed off-target homology-independent integrations, likely through a HITI-type insertion event. Efficient GFP expression as expected is only seen with the correct gRNA sequence and HDR mediated repair. Bars represent observed GFP+ percentages from T cells from one representative donor electroporated with the indicated components. **f,** Comparisons of on-target GFP expression vs functional off-target integrations across five templates reveal HDR is highly specific, but that off-target integrations can be observed at low frequencies. Average of two donors along with each donor’s exact value are graphed. **g,** A matrix of gRNAs and HDR templates were electroporated into CD4+ T cells from two healthy donors. The average GFP expression as a percentage of the maximum observed for a given template is displayed. Across six unique HDR templates and gRNAs, on-target HDR mediated integration was the by far most efficient. One HDR template, a C-terminal GFP fusion tag into the nuclear factor FBL, had consistently higher off-target expression across gRNAs, potentially due to a gene-trap effect as the 3’ homology arm for FBL contains a splice-site acceptor followed by the final exon of FBL leading into the GFP fusion.

**Extended Data Figure 11:**
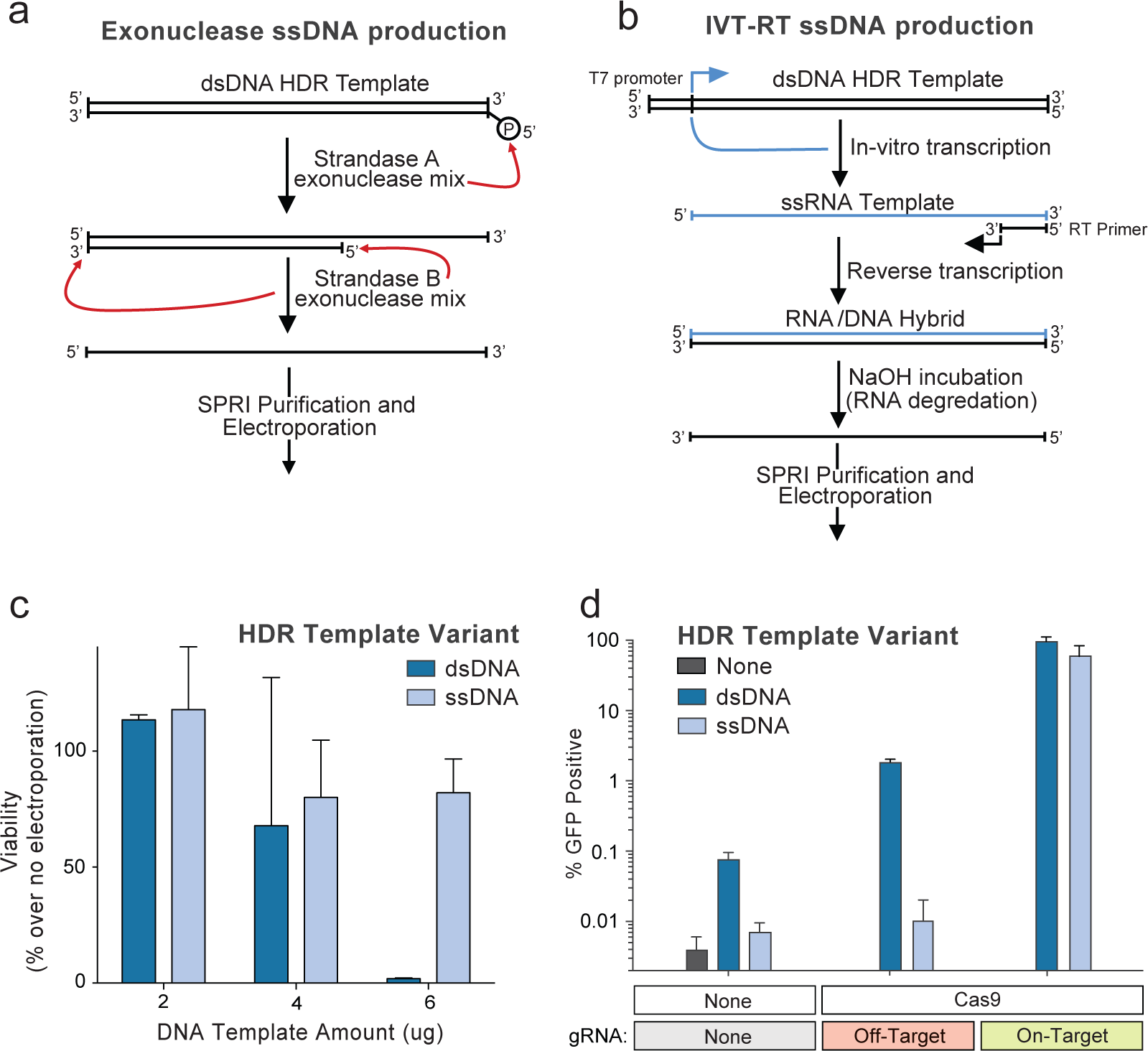
Production and usage of long ssDNA HDR templates. **a,** If a large enough amount of long ssDNA sequence could be produced for electroporation, off-target integrations could be reduced without compromising on-target efficiency. One method we developed involves a two-step selective exonuclease digestion that specifically degrades one strand of a PCR product that has been labelled by 5’ phosphorylation, easily added onto a PCR primer prior to amplification. **b,** We also applied a second ssDNA production method based on sequential *in vitro* transcription (IVT) and reverse transcription (RT) reaction. A PCR product with a short T7 promoter appended serves as an IVT template to produce a ssRNA product. Following annealing of an RT primer and reverse transcription, an RNA/DNA hybrid is formed which can be easily transformed into a long ssDNA template by incubation in sodium hydroxide, which selectively degrades the RNA strand. **c,** At 4 days post-electroporation, long ssDNA HDR templates did not show the decreasing viability in CD3+ T cells electroporated with a linear dsDNA HDR template of the same length. **d,** Electroporation of a ssDNA HDR template reduced off-target integrations to the limit of detection (comparable to levels seen with no template electroporated) both with no nuclease added and at induced off-target dsDNA breaks (Off-target gRNA + Cas9). Displayed data shows average and standard deviation of two healthy donors.

**Extended Data Figure 12:**
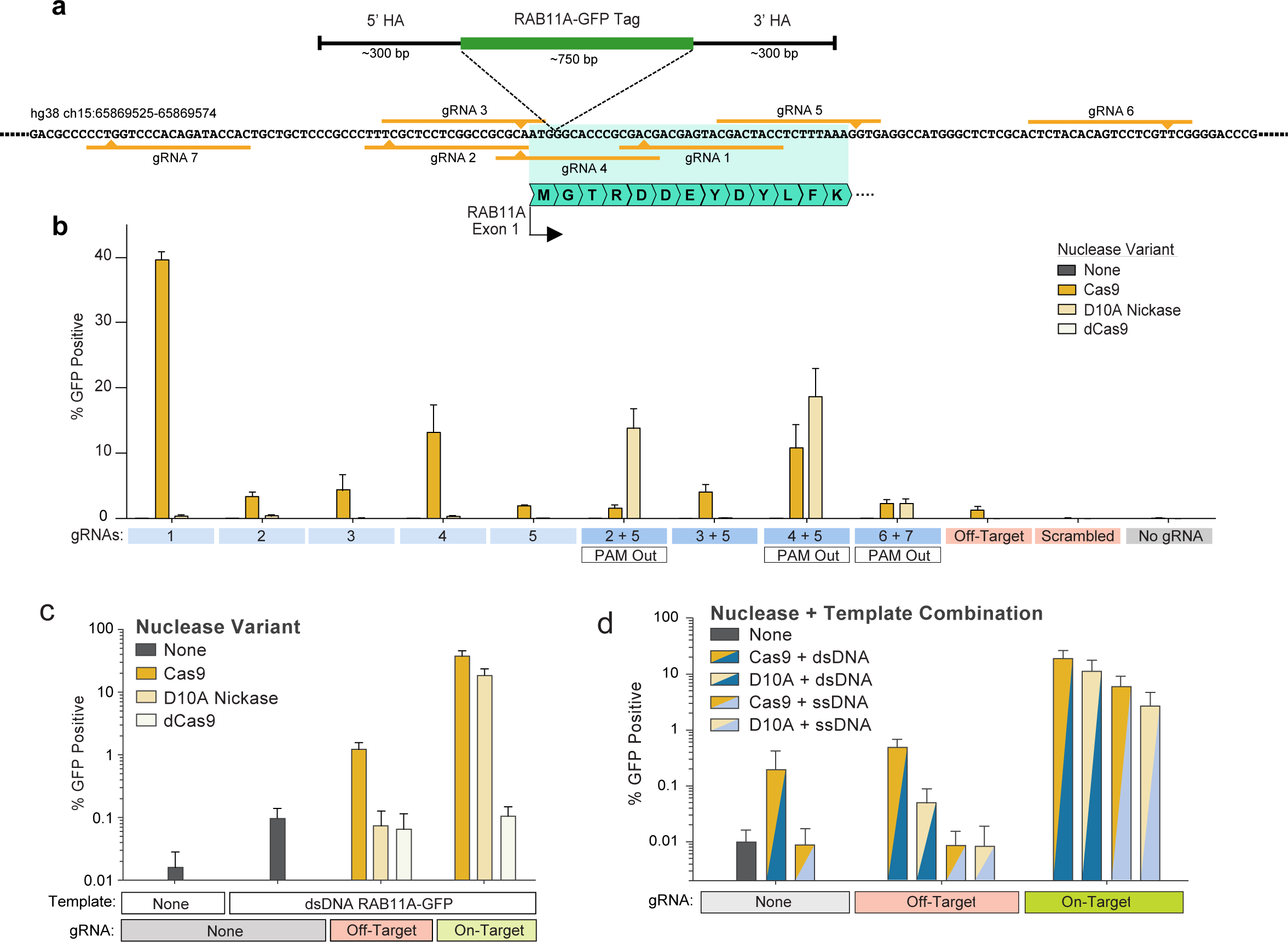
Efficient non-viral genome targeting with a Cas9 Nickase. **a,** Diagram of the genomic locus containing the first exon of *RAB11A.* Use of spCas9 with a single guide RNA (gRNA 1) along with a dsDNA HDR template integrating a GFP in frame with RAB11A directly after the start codon results in efficient GFP expression (**Fig. 1c**). Use of a Cas9 nickase (D10A variant) with two gRNAs reduces the incidence of off-target genome cutting. **b,** A series of individual gRNAs as well as dual gRNA combinations were tested for GFP insertion efficiency at the RAB11A N-terminal locus. As expected, no gRNAs showed appreciable levels of GFP insertion when using a nuclease dead Cas9 (dCas9). Multiple individual gRNAs cutting adjacent to the insertion site showed GFP integration when using Cas9, but none as efficiently as gRNA 1. The D10A nickase showed little to no GFP integration with individual guides, but with multiple two-guide combinations showed efficient GFP integration. Only in gRNA combinations where the two PAM sequences were directed away from each other (PAM Out) was GFP integration seen. **c,** GFP integration efficiencies as presented in (**b**) but graphed on a logarithmic scale reveal lower levels of functional off-target integrations ("off-target" gRNA, targeting CXCR4) when using the D10A nickase compared to spCas9, likely due to the requirement for the D10A nickase to have two gRNAs binding in close proximity to induce a dsDNA break. **d,** Long ssDNA templates could be successfully combined with Cas9 nickases (D10A) for targeted integration, similar to linear dsDNA templates. Long ssDNA HDR templates with D10A nickase showed lower efficiencies of GFP integration at the *RAB11A* site, but this appeared to be site specific, as at the combination of long ssDNA and D10A showed higher efficiencies compared to dsDNA and Cas9 at a second site (**Fig. 3d**). All graphs show the average and standard deviation from two healthy donors.

**Extended Data Figure 13:**
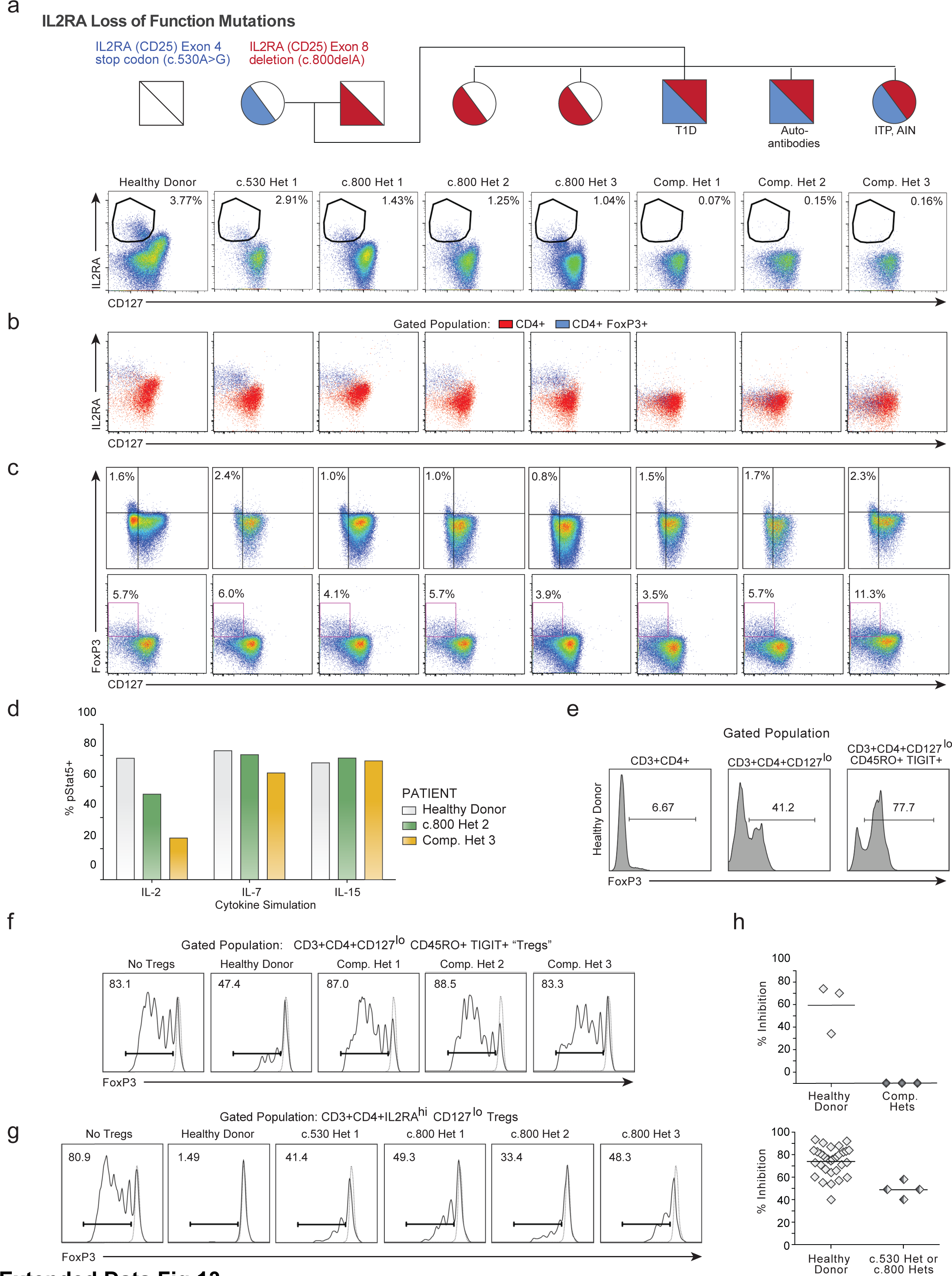
Reduced IL2RhiCD127lo Treg frequencies and defective Treg suppressive capacity in subjects with two loss of function IL2RA mutations. **a,** CD3+CD4+ T cells from a healthy donor and all family members, including *IL2RA* heterozygotes (c.530 het 1, c.800 hets 1-3) as well as compound heterozygotous children (Comp. Hets 1-3), with loss-of-function *IL2RA* mutations were analysed by flow cytometry to assess presence of IL2RAhiCD127lo Tregs. **b,** In healthy donors and single hets, CD4+FoxP3+ T cells are predominantly IL2RAhiCD127lo. In the compound heterozygotes, a CD127loCD4+FoxP3+ population is present, but does not express IL2RA. **c,** Clinical phenotyping performed at two separate sites confirms compound heterozygotes possess normal frequencies of CD127loFoxP3+ cells. **d,** Deficiency in IL2RA surface expression in compound heterozygote 3 led to aberrant downstream signalling as measured by phosphorylate (p)Stat5 expression after stimulation with IL-2, but not IL-7 or IL-15. **e,** Due to the inability to sort IL2RAhi Tregs from the IL2RA-deficient compound heterozygotes, an alternate gating strategy was established to enrich for FoxP3+ cells from CD3+CD4+ T cells using surface markers CD127loCD45RO+TIGIT+. Intracellular FoxP3 staining from the indicated gated population is shown. **f,** While these CD3+CD4+CD127loCD45RO+TIGIT+ potential "Tregs" were highly enriched for FoxP3 and showed some suppressive capacity when cultured with CFSE-labelled stimulated responder T cells (Tresps) from healthy donors, CD3+CD4+CD127loCD45RO+TIGIT+ from the compound heterozygotes showed no suppressive ability. Stimulated Tresp population (Solid curves), non-stimulated Tresp (Dashed curve). **g,** Correction of either IL2RA mutation in the compound heterozygotes individually would still leave the other mutation, leaving the cells as single heterozygotes. To confirm that such a potential correction would result in some level of functional suppression, we assessed the suppressive ability of CD4+IL2RAhiCD127lo Tregs from the c.530 and c.800 single heterozygote family members as in **(f)**. **h,** Dot plot summaries of Treg suppressive ability in cells from healthy donor, IL2RA-deficient compound heterozygotes **(f)** and IL2RA+/-c.530 or c.800 heterozygotes **(g)**. While CD3+CD4+CD127loCD45RO+TIGIT+ "Tregs" from compound heterozygotes showed no suppressive ability, conventional CD4+IL2RAhiCD127lo Tregs from the single heterozygote family members showed some suppressive capacity, consistent with their lack of pronounced clinical phenotype compared to the compound hets. Thus, correcting functional IL2RA expression on the surface of FOXP3+ T cells from these patients may represent a viable approach for developing an *ex vivo* gene therapy.

**Extended Data Figure 14:**
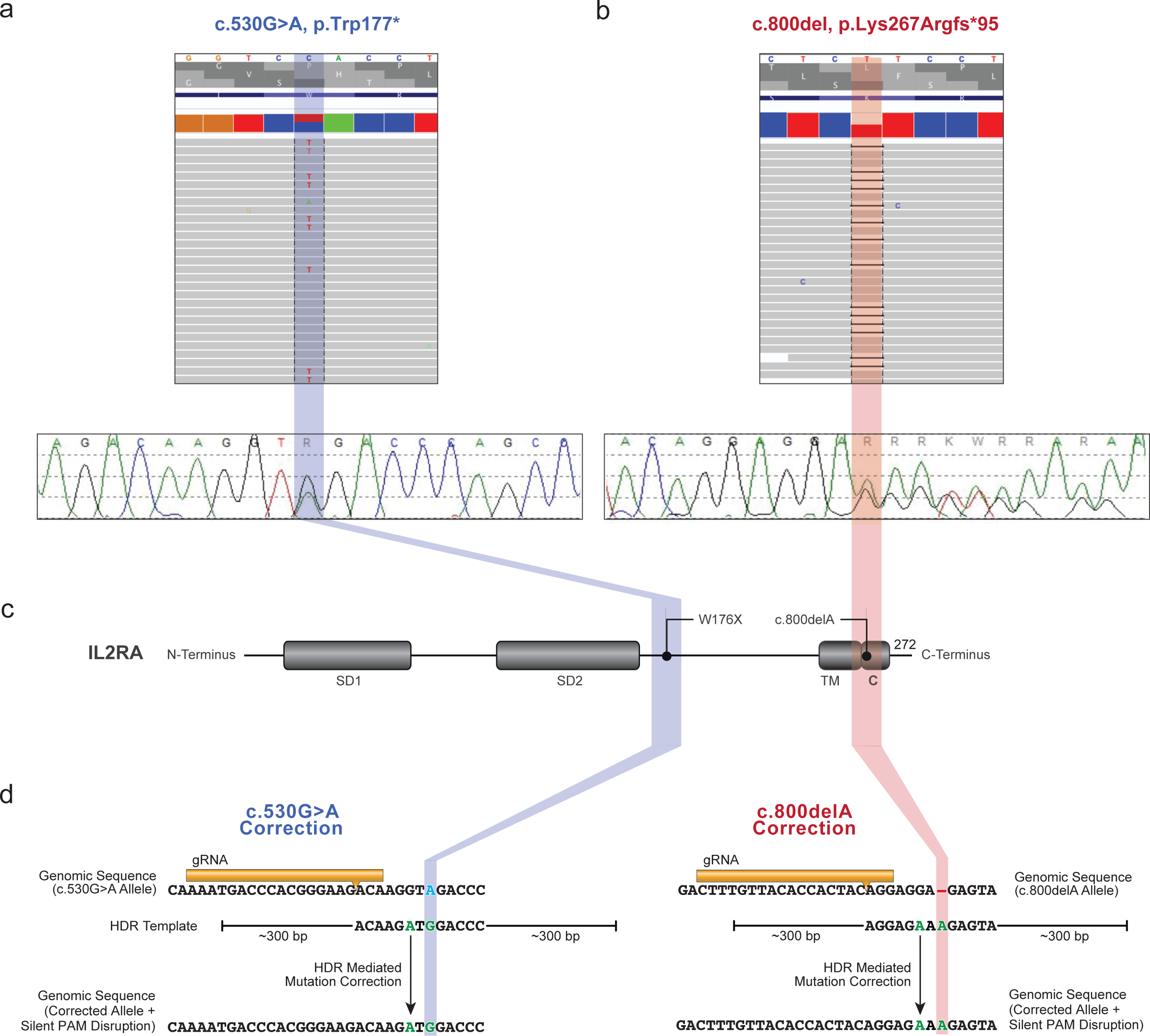
Identification of compound heterozygous mutations in IL2RA and design of corrective CRISPR-Cas9 genome targeting strategy. **a,** Initial genetic testing of the proband (**Supplementary Note 3**) using an in-house targeted next-generation sequencing multi-gene panel of over 40 genes known to be involved in monogenic forms of diabetes was negative. Subsequent exome sequencing in the trio of proband and parents revealed two causative mutations in the *IL2RA* gene. The mother possessed a single heterozygous mutation (c.530G>A) in exon 4 of *IL2RA,* resulting in a premature stop codon. **b,** The father possessed a single heterozygous mutation (c.800delA) in exon 8 of IL2RA, resulting in a frameshift mutation leading to a 95 amino acid long run-on. Sanger sequencing confirmed that the proband was a compound heterozygote for both mutations. **c,** A linear depiction of the IL2RA protein annotated with approximate locations of the two identified IL2RA mutations. Annotated functional domains are shown: SD1, sushi domain 1; SD2, sushi domain 2; TM, transmembrane; C, cytoplasmic. **d,** The genomic sequences including the specified mutations were used to design CRISPR-Cas9 genome targeting reagents to correct the two *IL2RA* mutations. A gRNA was designed to cut adjacent to the site of each mutation, 8 bps away for c.530 mutation (Blue), and 7 bps away for c.800 (Red). For each mutation, an HDR template was designed including the corrected sequence (Green) as well as a silent mutation in a degenerate base to disrupt the PAM sequence ("NGG") for each guide RNA. Displayed genomic regions (not to scale) for c.530 mutation site (hg38 ch10:6021526-6021557) and c800 mutation site (hg38 ch10:6012886-6012917). Both ssODN HDR Templates (ssDNA with 60 bp homology arms), and large dsDNA or ssDNA HDR Templates (as displayed, with ~300 bp homology arms) were used.

**Extended Data Figure 15:**
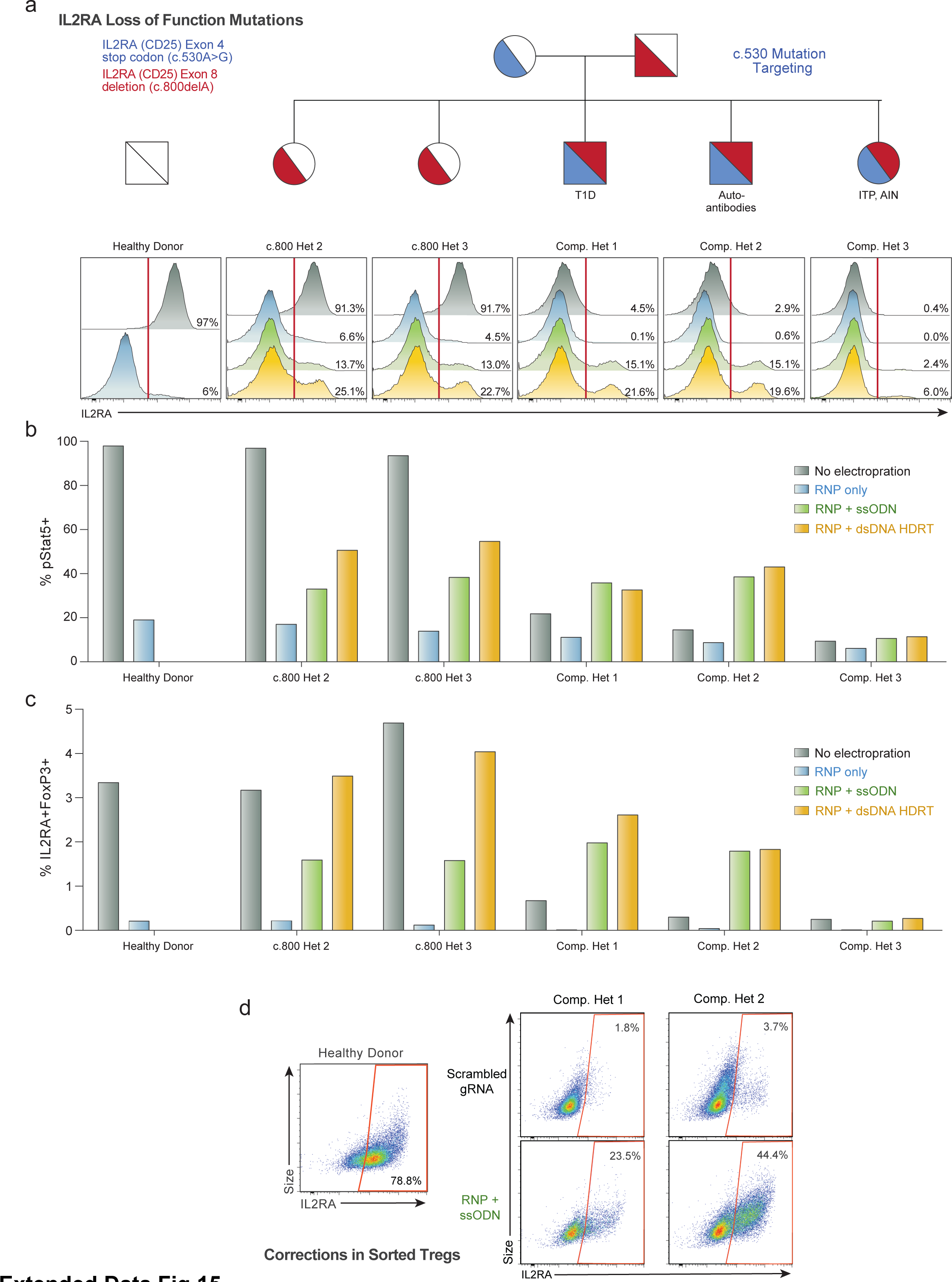
HDR-mediated correction of IL2RA c.530A>G loss of function mutation. **a,** Unlike the gRNA targeting the c.800delA mutation at the C-terminus of IL2RA **(Extended Data Fig. 16)**, the gRNA targeting the c.530A>G mutation (causing a stop codon in an interior exon) results in substantial (~90%) loss of IL2RA cell surface expression in a healthy donor and single heterozygotes (c.800 Het 2 and 3) 2 days following electroporation of the RNP alone (Blue) into CD3+ T cells. While starting from a very small IL2RA+ percentage, this reduction was also observed in all three compound heterozygotes, potentially as some small amount of protein can be surface expressed off of the c.800delA allele. This reduced IL2RA expression could be partially rescued by inclusion of an ssODN HDR template (Green) and even more substantially rescued using a large dsDNA HDR template (Yellow). Both template types contained the corrected sequence, a silent mutation to remove the gRNA’s PAM sequence, and either 60 bp (ssODNs) or ~300 bp (large dsDNA) homology arms (Yellow) (**Extended Data Fig. 14**). Unlike targeting of the c.800delA mutation for correction, increased IL2RA surface expression in T cells from the compound heterozygotes is only seen when an HDR template is included. In all three compound heterozygotes, the dsDNA HDR template yielded greater percentages of IL2RA+ cells. **b,** Increased phosphorylated (p)Stat5 in response to IL-2 stimulation (200 U/mL) 7 days following electroporation in CD3+ T cells from compound heterozygote patients undergoing HDR-mediated mutation correction compared to no electroporation or RNP only controls. pStat5+ cells correlated with increases in IL2RA surface expression. **c,** Similarly, increased proportions of IL2RA+FoxP3+ cells are seen 9 days following electroporation in the HDR correction conditions in compound heterozygote patients. Lower percentages of correction were seen when targeting the c.530 mutation for HDR correction in compound heterozygote 3, potentially due altered cell-state associated with the patient’s disease or the patient’s immunosuppressive drug regimen (**Extended Data Fig. 17**). **d,** Mutation correction was possible in sorted Treg-like cells from the affected patients. CD3+CD4+CD127loCD45RO+TIGIT+ "Tregs", a population highly enriched for FoxP3+ cells (**Extended Data Fig. 13**) identified without the traditional Treg IL2RA surface maker (absent due to the causative mutations), were FACS sorted and underwent correction of the c.530A>G mutation using a Cas9 nuclease and short ssDNA HDR template (ssODN). After 12 days in culture, during which time the cells expanded >100 fold, greater than 20% (compound het 1) and 40% (compound het 2) of targeted cells expressed IL2RA on their surface, demonstrating functional correction and expansion of a therapeutically relevant cell type. All electroporations were performed according to optimized non-viral genome targeting protocol (**Methods**). For ssODN electroporations, 100 pmols in 1uL H2O were electroporated.

**Extended Data Figure 16:**
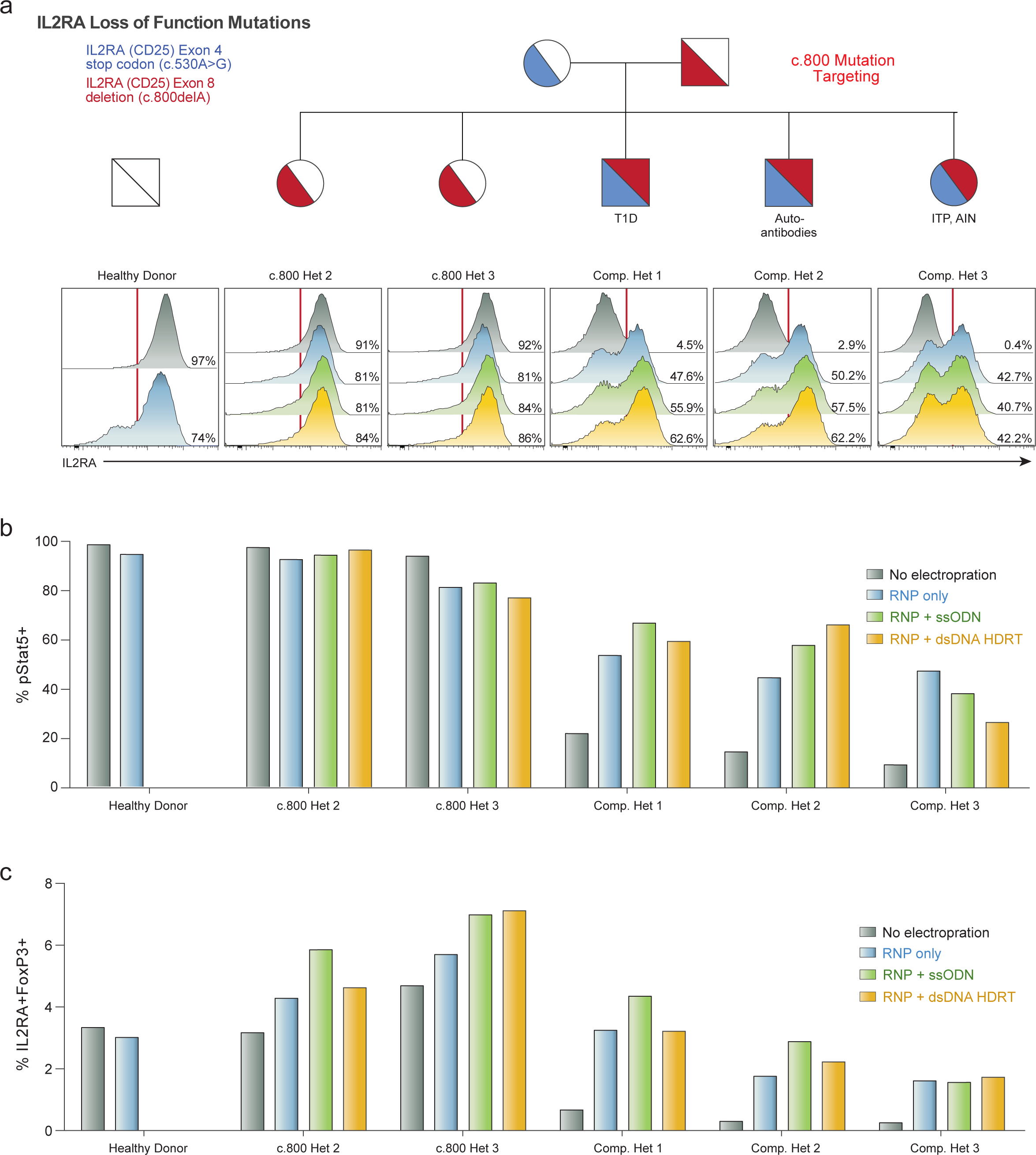
HDR and Non-HDR-mediated correction of *IL2RA* c.**800delA frameshift loss-of-function mutation.** **a,** Histograms of IL2RA surface expression in CD3+ T cells in all children from a family carrying two loss-of-function *IL2RA* mutations, including three compound heterozygotes that express minimal amounts of IL2RA on their surface (No electroporation, Grey). Two days following electroporation of an RNP containing a gRNA against the site of one of the two mutations, a one base pair deletion in the final exon of *IL2RA* (c.800delA) causing a run-on past the normal stop codon, CD3+ T cells from a healthy donor and single hets (c.800 Het 2 and 3) showed slight increases in IL2RA-cells (RNP only, Blue). This modest reduction is potentially due to the gRNA targeting the C-terminus of the protein where small indels may cause less pronounced loss of surface protein expression. Surprisingly, the RNP alone resulted in IL2RA surface expression in almost 50% of edited T cells in all three compound heterozygotes. In cells from two of the compound heterozygous children, increases in the percent of cells with IL2RA correction compared to RNP only could be achieved by inclusion of an ssODN HDR template sequence with the mutation correction (RNP+ssODN, Green), and further increased when using a longer dsDNA HDR template to correct the mutation (RNP + dsDNA HDRT, Yellow) **(Extended Data Fig. 14)**. **b,** Stat5 phosphorylation (pStat5) in response to high dose IL-2 stimulation (200 U/mL) in edited CD3+ T cells following 7 days of expansion post-electroporation. Increased numbers of pStat5+ cells correlated with increases in IL2RA surface expression **(a)**. **c,** Following 9 days of expansion postelectroporation, intracellular FoxP3 staining revealed a marked increased proportion of IL2RA+ FoxP3+ cells in CD3+ T cells compared to no electroporation controls, approaching the proportion of IL2RA+ FoxP3+ cells seen in similarly cultured cells from a healthy donor. Electroporations were performed according to optimized non-viral genome targeting protocol (**Methods**). For ssODN electroporations, 100 pmols in 1uL H2O were electroporated.

**Extended Data Figure 17:**
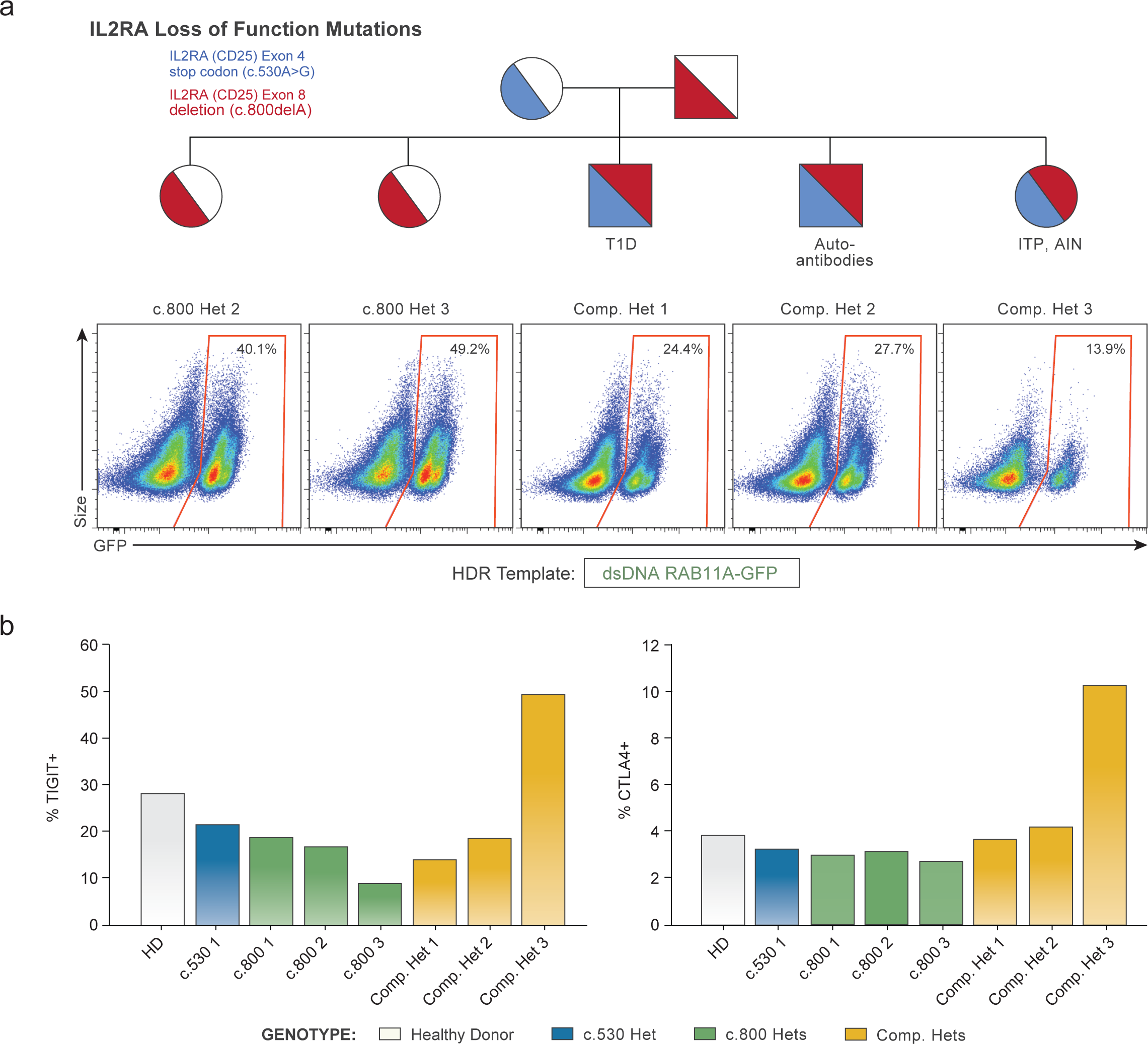
Diminished HDR potential and altered clinical phenotype in IL2RA loss-of-function patient receiving immunosuppressants. **a,** Flow cytometric analysis of GFP expression 6 days following electroporation of a positive HDR control RAB11A-GFP dsDNA HDR template into CD3+ T cells from the indicated patients revealed lower GFP expression in the three compound heterozygotes compared to their two c.800 heterozygote siblings. Compared to a cohort of twelve healthy donors similarly edited **(Fig. 1d)**, both c.800 heterozygotes as well as compound het 1 and 2 were within the general range observed across healthy donors, whereas compound het 3 had lower GFP expression than any healthy donor analysed. Of note, in compound het 3 HDR-mediated correction at the c.530 mutation was substantially lower than the other two compound hets **(Fig. 3a)**, IL2RA surface expression after electroporation of the c.800delA targeting RNP alone was similar **(Extended Data Fig. 16)**. Unlike HDR-mediated repair, a NHEJ mediated frameshift correction at c.800delA may not require cell proliferation, consistent with compound het 3 being the only compound heterozygote patient on active immunosuppressants at the time of blood draw and T cell isolation **(Supplementary Note 3)**. **b,** Altered cell-state associated with the patient’s disease could also be contributory to diminished HDR rates. TIGIT and CTLA4 expression levels in non-edited, isolated CD4+ T cells from each indicated patient measured by flow cytometry. Consistent with altered cell states and or/ cell populations, cells from compound het 3 had a distinct phenotype, with increased TIGIT and CTLA4 expression compared both to healthy donors, the heterozygous family members, as well the other two compound heterozygous siblings.

**Extended Data Figure 18:**
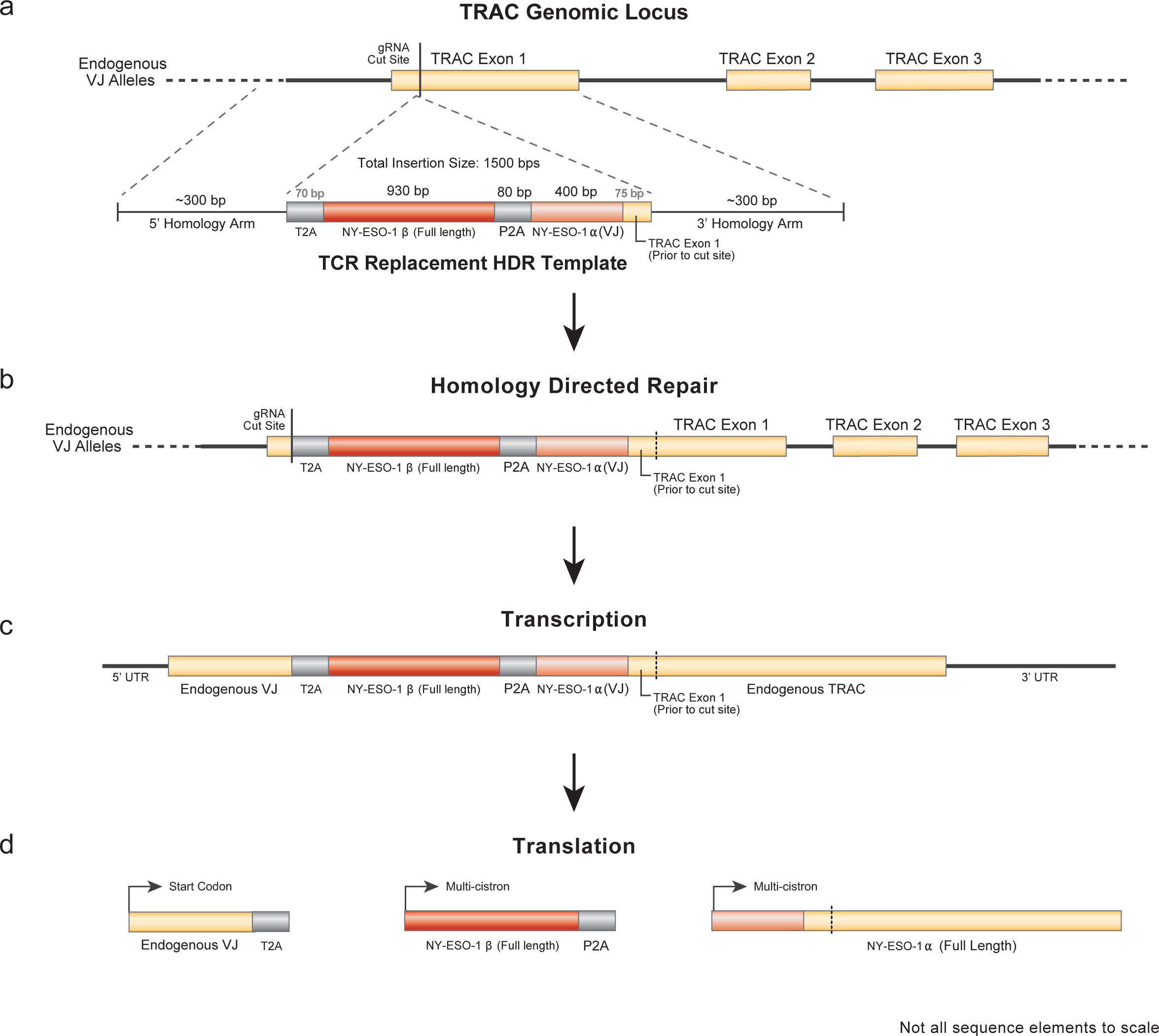
Diagram of Endogenous TCR Replacement With a NonViral HDR Template. **a,** The genomic locus of the T cell receptor is extremely complicated, with a large variety of variable alleles (V and J alleles for the TCR-α chain and V, D and J alleles for the TCR-β chain) that undergo somatic gene rearrangement during T cell development in order to produce a functional T cell receptor. Important for the diversity of the TCR repertoire but challenging for targeted genomic editing at the TCR locus (whether knockouts or knock-ins), these recombined sequences are variable across the polyclonal population of T cells. However, for both the TCR-α and TCR-ß chains there is a constant domain at the C-terminus of the protein that is shared by all T cells, no matter what V-J or V-D-J segments have been rearranged. This constant sequence, termed T Receptor Alpha Constant *(TRAC)* Exons 1, Exon 2, etc. allows for a single gRNA sequence to be used to modify every T cell no matter what rearranged TCR they express. **b,** A 2.1 kb HDR Template was used to replace the endogenous TCR. Approximately 300 bp homology arms surround a ~1.5 kb inserted sequence encoding a self-excision peptide followed by the full-length sequence of the TCR-β chain of the desired antigen specific T cell (here the 1G4 NY-ESO-1 specific TCR). A second self-excision peptide follows the TCR-β chain, and separates it from the variable (recombined V and J) sequence of the desired antigen specific TCR-α chain. **c,** Only the variable region from the TCR-α chain and the sequence of the TRAC exon 1 prior to the gRNA cut site needs to be inserted, as the remaining TRAC exons are spliced together into the final mRNA sequence. **d,** This targeting strategy is designed to yield three peptide chains: a remnant endogenous variable region peptide that does not possess a transmembrane pass (and thus should be degraded in the endoplasmic reticulum or secreted following transcription), the full-length desired antigen specific TCR-ß chain, and the full-length desired antigen specific TCR-α chain. The result is a T cell that expresses both chains of a desired antigen specific TCR under the control of the endogenous TCR-α promoter.

**Extended Data Figure 19:**
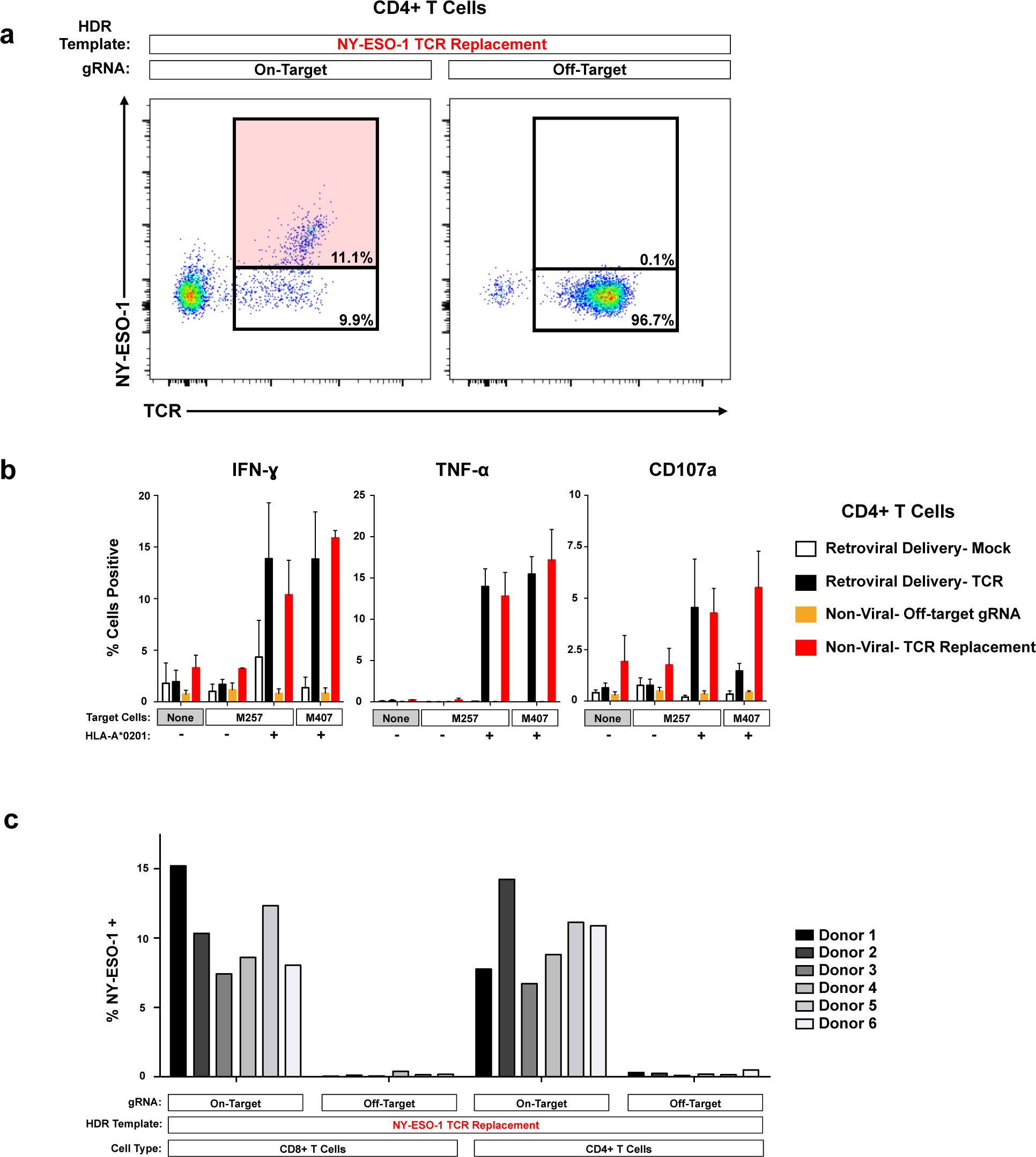
Reproducibility of TCR Replacement Across Donors and in CD4 and CD8 T Cells. **a,** Non-viral replacement of the endogenous TCR in CD4+ T cells from the same donor displayed in **Fig. 4b**. **b,** Functional cytokine production is observed selectively following antigen exposure in targeted CD4+ T cells, similarly to CD8+ T cells (**Fig. 4c**). Graphs display average and standard deviation across two healthy donors. **c,** Non-viral TCR replacement was consistently observed in both CD8+ and CD4+ T cells across a cohort of six healthy blood donors.

**Extended Data Figure 20:**
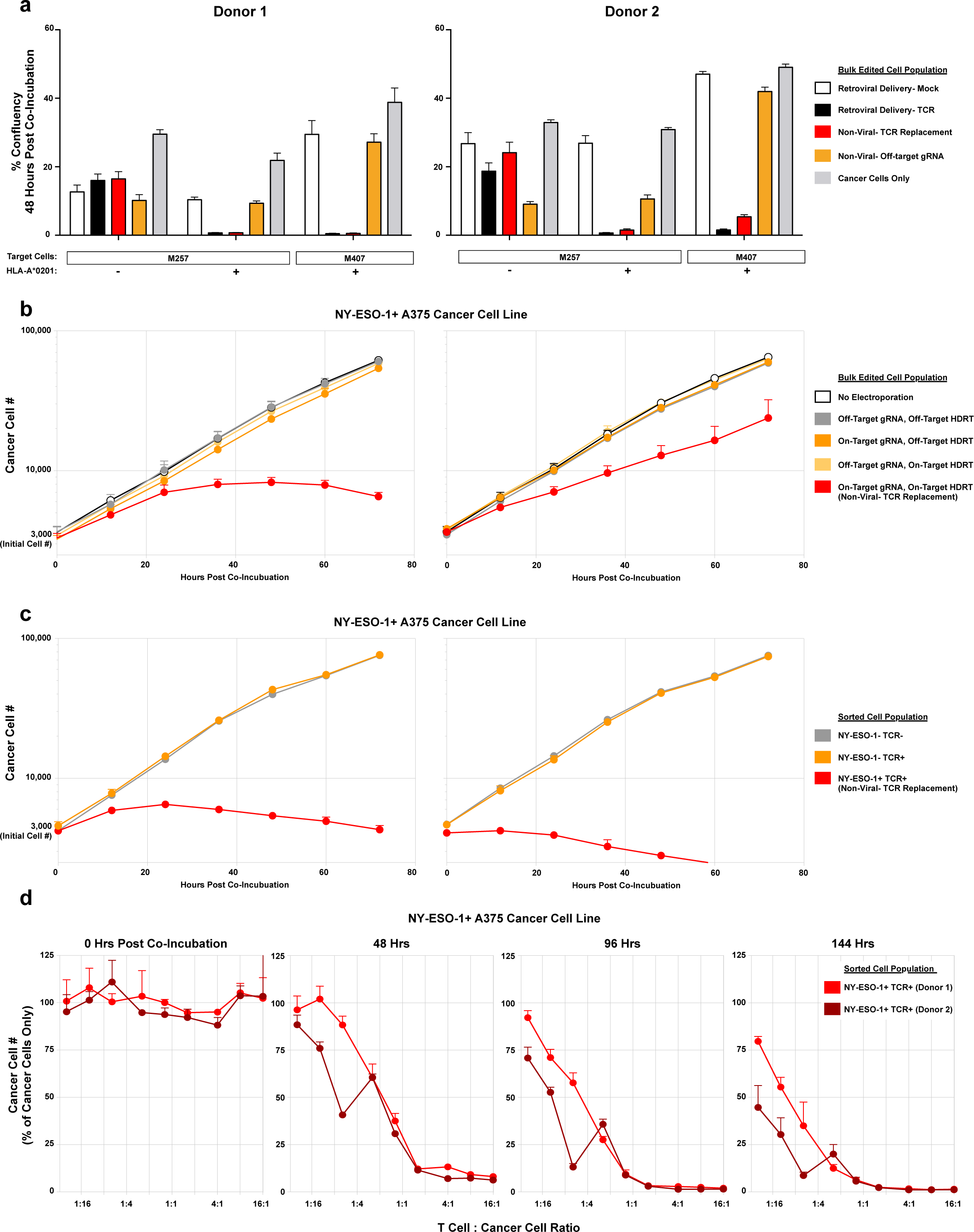
Antigen-Specific Killing by T Cells with a Non-Virally Replaced TCR. **a,** Indicated melanoma cell lines were co-incubated with the indicated bulk-edited T cell populations at a ratio of 5:1 T cells to cancer cells. At 48 hours post co-incubation the percent cancer cell confluency was recorded with by automated microscopy (where nuclear RFP marks the cancer cells). T Cells expressing the NY-ESO-1 antigen specific TCR, either by retroviral transduction (Black) or by non-viral knock-in endogenous TCR replacement (Red) both showed robust target cell killing only in the target cancer cell lines expressing NY-ESO-1 and the matched HLA-A*0201 class I MHC allele. **b,** To ensure that target cell killing by non-viral TCR replacement T cells (Red) was not due to the either the gRNA or the HDR template used for TCR replacement alone, a matrix of on/off target gRNAs and on/off target HDR templates was assayed for target cell killing of the NY-ESO-1+ HLA-A*0201 + A375 cancer cell line (off-target gRNA and HDRT were specific for RAB11A-GFP fusion protein knock-in). Only cells with both the on-target gRNA as well as the on-target HDR template demonstrated target cell killing. **c,** Target cell killing by non-viral TCR replacement T cells was due specifically to the NY-ESO-1-recognizing TCR+ cell population observed by flow cytometry after non-viral TCR replacement (**Fig. 4b**). Starting with the bulk edited T cell population (all of which had been electroporated with the on-target gRNA and HDR template), we separately sorted three populations of cells: the NY-ESO-1+TCR+ cells (non-virally replaced TCR) (red), the NY-ESO-1-TCR-cells (TCR knockout) (grey), and the NY-ESO-1-TCR+ cells (those that retained their native TCR but did not have the NY-ESO specific knock-in TCR) (orange). Only the sorted NY-ESO-1+ TCR+ population demonstrated target cell killing (4:1 T cell to cancer cell ratio). **d,** Sorted NY-ESO-1 + TCR+ cells from a bulk T cell edited population (on-target gRNA, on-target HDR template) showed a strong dose-response effect for target cancer cell killing. Within 48 hours T cell to cancer cell ratios of 2:1 and greater showed almost complete killing of the target cancer cells. By 144 hours, even T cell to cancer cell ratios of less than 1:16 showed evidence of robust target cell killing. All graphs show average and standard deviation from three technical replicates per displayed condition/time-point.

**Extended Data Table 1:**
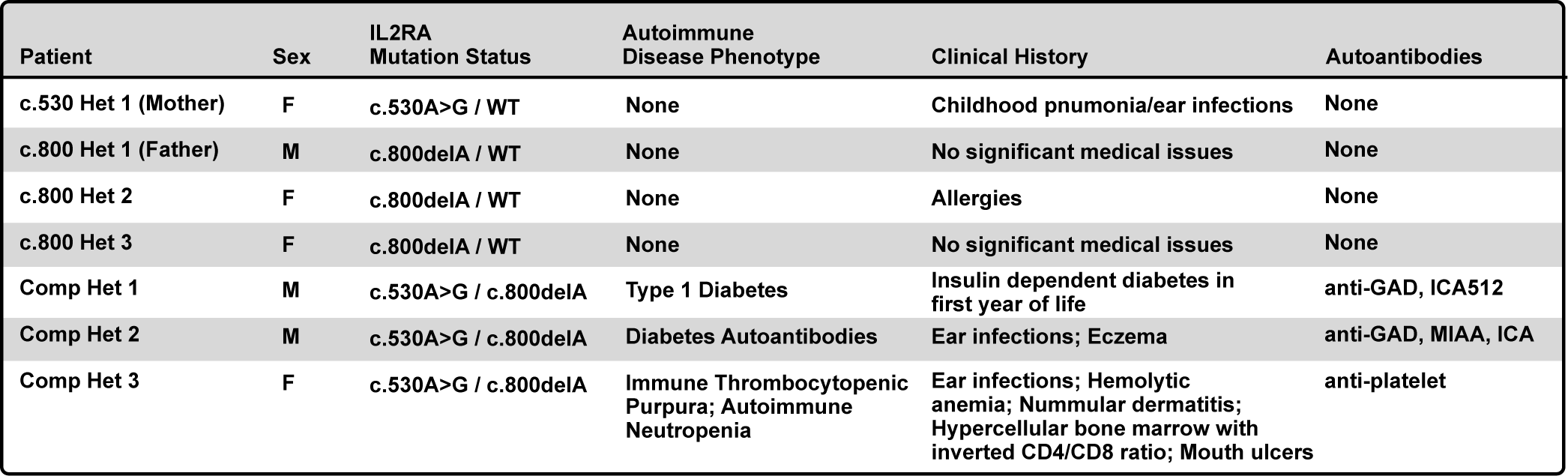
Clinical history of family members with *IL2RA* mutations. Mutation status and clinical phenotypes of members of a family with two distinct *IL2RA* coding mutations, including three patients with compound heterozygote mutations.

## SUPPLEMENTARY INFORMATION

1. **Reporting Summary**
2. **Supplementary Table 1** A list of antibodies used in this study
3. **Supplementary Table 2** A list of HDR template, DNA primer, and gRNA sequences used in study.
4. **Supplementary Note 1** Heterozygous/Homozygous integration prediction model.
5. **Supplementary Note 2** Analysis of off-target effects
6. **Supplementary Note 3** Clinical history, genetic testing, and clinical phenotyping of family with compound heterozygous *IL2RA* mutations.

### Supplementary Note 1: Heterozygous/Homozygous integration prediction model

An estimation of the percentage of cells with bi-allelic insertions at a single autosomal genomic locus (two potential alleles) can be made from only fluorescent phenotypes if two HDR templates integrating different fluorescent proteins into that same site are introduced into the cell (by electroporation). A simple probability model requires only two assumptions.

*Assumption 1:* There are no off-target integrations at other sites besides the target locus that contribute to fluorescent phenotypes.

*Assumption 2:* Integration of a specific second fluorescent protein (i.e. RFP) does not depend on which fluorescent protein was integrated at the cell’s other allele (i.e. GFP or RFP integrations one the first allele are equally likely to have an RFP integration at the second).

Following the labelling in fig. S13A-C, the percentages of four different phenotypic populations are known:

- % GFP^−^RFP^−^
- % GFP^+^RFP^−^
- % GFP^−^RFP^+^
- % GFP^+^RFP^+^

From these, immediately two genotypes are known:

1) Genotype A = NA/NA = % GFP^−^RFP^−^
2) Genotype E = GFP/RFP = % GFP^+^RFP^+^

The four remaining genotypes sum to the two remaining single fluor positive phenotypes:

3) Genotype B + Genotype D = GFP/NA + GFP/GFP = % GFP^+^RFP^−^
4) Genotype C + Genotype F = RFP/NA + RFP/RFP = % GFP^−^RFP^+^

The probabilities that a RFP^+^ cell will also be GFP^+^, and vice versa, are also known from the phenotypes:

5) Probability of being GFP^+^ given being RFP^+^ = P(GFP|RFP) = (% GFP^+^RFP^+^) / (% RFP^+^ + %GFP^+^RFP^+^)
6) Probability of being RFP^+^ given being GFP^+^ = P(RFP|GFP) = (% GFP^+^RFP^+^) / (% GFP^+^ + %GFP^+^RFP^+^)

Following from assumption 2, if the probability that a cell receives a GFP integration at its second allele is independent of whether the first integration was a GFP or RFP, then a relationship between the single positive genotypes can be determined (fig S13):

7) D = P(GFP|RFP) * B
8) F = P(RFP|GFP) * C

Inserting the equations 7 and 8 into equations 3 and4 respectively and simplifying solves for the remaining genotypes in terms of the known phenotypes:

9) B = % GFP^+^RFP^−^ / (1 + (% GFP^+^RFP^+^) / (% RFP^+^ + %GFP^+^RFP^+^))
10) C = % GFP^−^RFP^+^ / (1 + (% GFP^+^RFP^+^) / (% GFP^+^ + %GFP^+^RFP^+^))
11) D = % GFP^+^RFP^−^−B
12) F = % GFP^−^RFP^+^−C

From the known genotypes, the observed % of cells that are have mono-allelic or bi-allelic insertions, as well as other statistics, can be calculated readily:

- Observed % Cells Heterozygous = B + C
- Observed % Cells Homozygous = D + E + F
- Observed % Cells with at least 1 insertion = B +C+ D + E +F=1 -A = 1 -%GFP-RFP-
- Observed % Alleles that have a GFP = (B + E + 2D) / 2
- Observed % Alleles that have a RNP = (C + E + 2F) / 2
- Observed % Alleles with an insertion = % Alleles_GFP_ + % Alleles_RFP_

An expected % of cells homozygous if the HDR alleles were distributed randomly (in essence at Hardy-Weinberg Equilibrium) can be calculated from the observed % of cells with at least one insertion (HDR):

- p = HDR allele (GFP or RFP)
- q = non-HDR allele (NA)
- X = % of cells observed to have at least one HDR

13) p + q = 1
14) p^2^ + 2*p*q + q^2^ = 1

As any cell that has an HDR (GFP or RFP) allele will show the phenotype (in this case GFP+ or RFP+):

15) X = p^2^ + 2*p*q

Substituting X into equation 14 and simplifying:

16) q = (1 - X)^1/2^
17) p = 1 - q
18) p = 1 - (1 -X)^/2^

p^2^ will give then give the expected % of cells homozygous for HDR integration if HDR template insertion was random among the target alleles:

19) p^2^ = 2 − 2(1 − X)^1/2^ − X

As X is known, the expected % of homozygous cells can be calculated directly from the observed total % of cells with at least one HDR, and can then be compared the observed % of homozygous cells calculated by taking into account the information provided by integration of two separate fluorophores.

### Supplementary Note 2: Analysis of off target effects

A common concern for use of genetically modified T cells therapeutically is the potential for off-target effects. With any targeted editing strategy, there are at least three potential types of undesirable outcomes. First, the nuclease (such as a Cas9 RNP) can generate non-specific mutations via-nonhomologous repair mechanisms at the cleavage site or at off-target sites. Second, the DNA HDR template (either viral or non-viral) could integrate off-target either at off-target Cas9 RNP cut sites or at naturally occurring double-stranded breaks in host DNA^22^ (**Extended Data Fig. 10a, b**). Even integrase-deficient AAVs templates have been shown to stably integrate at off-target sites^8, 21^. Third, the DNA HDR template could integrate at the desired target site but integrate incorrectly, for example through homology-independent mechanisms^23^. Off-target cutting and integrations must be minimized in cells destined for therapeutic use to ensure that integrated sequences remain under proper endogenous regulation and that critical offtarget sites are not disrupted.

We looked for unintended non-homologous integrations with the non-viral system using an N-terminal GFP-RAB11A fusion construct that contained the endogenous *RAB11A* promoter sequence within its 5’ homology arm. This construct can express GFP at off-target integration sites, which allowed us to assay for off-target events using flow cytometry (**Extended Data Fig. 10a**). Inclusion of a gRNA designed to cut a genome region that is not the homologous region to the targeting sequence can be used to measure integration at an off-target cut site. While efficient GFP expression depended on pairing the HDR template with the correct gRNA targeting that site, rare GFP+ cells (~0.1%) were observed when dsDNA HDR templates were delivered either alone or with an "off target" Cas9 RNPs (**Extended Data Fig. 10c-g**),

We tested whether nonviral genome targeting is compatible with a D10A Cas9 nickase variant engineered to reduce the potential for off-target double strand breaks^26, 27^. This variant requires that two gRNAs bind and cleave in close proximity to each other to produce a double strand break, thus reducing the number of off-target double stranded DNA breaks because both gRNAs would need to have an off-target cut site in close proximity to each other. We tested a series of gRNA combinations at the *RAB11A* locus for the integration of GFP, and found that guides in a “PAM-Out” orientation led to the efficient introduction of GFP when the D10A nickase was used (**Extended Data Fig. 12a, b**). The D10A nickase also reduced integrations at off-target cut sites, which we modelled using a gRNA that does not cut at a site homologous to the template. With this single “off-target” Cas9 nickase RNP, GFP integrations occurred only at background levels (**Extended Data Fig. 12c**).

While the D10A nickase may be useful to reduce off-target integration, a small, but significant number of GFP+ cells were generated even without targeting Cas9 with a gRNA to the site of homology. GFP+ cells were found at a similar rate when the donor along was used without any Cas9 nuclease, perhaps resulting from integration at naturally occurring double stranded DNA breaks (**Extended Data Fig. 10e**). We reasoned that the remaining off-target integrations could be eliminated by replacing the dsDNA HDR templates with similar ssDNA HDR templates, which are unable to integrate non-specifically at double strand breaks^24, 25^. To test this hypothesis, we generated ssDNA HDR templates using two methods we recently developed to produce the large amounts of long ssDNA required for electroporation^25^ (**Extended Data Fig. 10a, b**). ssDNA HDR templates reduced the number of functional off-target integrations approximately 100-fold, while maintaining efficient on-target integration (**Extended Data Fig. 10c, d**).

We next used a deep sequencing approach to assess off-target integrations and incorrect/non-homologous integration events that occurred with dsDNA and ssDNA HDR templates. We performed targeted locus amplification (TLA) sequencing in two donors with both dsDNA and ssDNA HDR templates (**Extended Data Fig. 9a**). Targeted locus amplifications were used due to the large amount of HDR template DNA that was retained in an episomal state in the cells. With both dsDNA and ssDNA HDR templates, no off-target integration sites were found above the limit of detection (approximately 1% of alleles) (**Extended Data Fig. 9b**). However, sequencing of the on-target locus revealed that some incorrect on-target integration events were detected with a dsDNA HDR template. One incorrect homology directed repair sequence was detected, resulting from 9 bps of overlap between a portion of GFP’s sequence and a region within the 3’ homology arm, which of course is also present at the target genomic locus. The ssDNA HDR template showed 10-fold fewer incorrect HDR integrations (**Extended Data Fig. 9c**). Where potential off-target cutting and/or integration activity is a concern, successful targeting can be achieved by combining D10A Cas9 nickase and ssDNA HDR templates.

### Supplementary Note 3: Clinical history, genetic testing, and clinical phenotyping of family with compound heterozygous *IL2RA* mutations

#### Clinical History of Family with Autoimmunity/Immune Dysregulation

The proband is a Caucasian infant who presented at 15 weeks of age after vomiting, fussiness and tachypnea led to medical evaluation that revealed severe diabetic ketoacidosis and a serum glucose level of 920 mg/dL. One week after diagnosis, testing for GAD65, IA-2 and insulin autoantibodies was negative; however, autoimmune diabetes was confirmed when repeat antibody tests at 5-7 months of age in three different laboratories showed positive results for IA-2 and insulin autoantibodies, as well as very high levels of GAD65 antibodies in two of the laboratories [42.8 nmol/L (<0.02) at Mayo Laboratories and 896 IU/mL (0.0-5.0) at Barbara Davis Center]. Testing for thyroid dysfunction and celiac disease has been negative but mildly low IgA levels suggest partial IgA deficiency. C-peptide testing was repeatedly completely undetectable, including at 7 months of age when measured 90 minutes after a feed with a serum glucose level of 202 mg/dL, at which time proinsulin was also undetectable. After the initial DKA was treated with intravenous insulin, the patient was discharged on multiple daily injections of subcutaneous insulin (glargine and lispro) initially and later transitioned to an insulin pump with continuous glucose monitoring. He consistently required a high replacement dose of insulin in the range of 0.8-0.9 units/kg/day (48% basal at 7 months of age). He had been delivered by repeat c-section at 37 weeks gestation with a birth weight of 3.629 kg (75th percentile) without any complications and there have been no concerns about his developmental progress and his medical history has otherwise been unremarkable. His parents have disparate Caucasian ancestry and denied consanguinity.

Clinical information on family members is provided in **Extended Data Table 1**. More detailed information is as follows:

1. Mother (37):
  a. Pneumonia as a child -explained as viral
  b. Ear infections as a child treated with antibiotics
  c. Tooth problems (perhaps related to antibiotics)
  d. Her father developed insulin dependent diabetes in his 30’s. He had a low WBC and also had nummular dermatitis of the scalp.
  e. Her mother had lupus
2. Father (44)
  a. Moroccan descent
  b. No major medical problems
  c. Some possible concern this his response time to common viral infections may be prolonged.
3. Affected child (14)
  a. Immune thrombocytopenic purpura: (+ anti-platelet antibodies)
  b. Neutropenia (anti-neutrophil Ab)
  c. Autoimmune haemolytic anaemia (DAT+ i.e. direct Coombs+)
  d. Nummular dermatitis of the scalp
  e. Hypercellular bone marrow: inverted CD4/CD8 ratio (0.36).
  f. Mouth ulcers
  g. Ear infections treated with tubes
  h. Diarrhoea as a child
  i. 46XX -no known chromosomal abnormality
  j. Flow cytometry of peripheral blood: 82.7% of CD45+ cells are CD3+ and 5.9% are CD19+. CD19+CD5+ cells are the deficient B cells. 43.6% of CD45+ cells are CD8+ with an inverted CD4/CD8 ratio (0.6). There is a relative increase in TCR(alpha beta)+ CD3+ CD4-CD8-T lymphocytes (26% of TCR alpha beta+ CD3+ cells and 5% of CD45+ leukocytes).
  k. Has been treated with immunosuppression including prednisone (20 mg), IgG-pro-IgA, Flonase nasal spray and topical steroids and Symbicort. Also treated with Neupogen.
4. Affected child
  a. 3+ diabetes autoantibodies (anti-GAD, MIAA, ICA, negative ZnT8 and ICA512/IA-2) normal OGTT
  b. Ear infections treated with tubes at 1 year
  c. Eczema in the winter
5. Unaffected daughter (15)
  a. Allergies, but otherwise healthy
6. Affected son (4)
  a. Eczema in winter
  b. b. Positive test for HSV
  c. c. Insulin dependent diabetes within the first year of life, C-peptide <0.1 at presentation, anti-GAD ab+ (>30 (nl<1U/ml) 1 yr after dx but negative at dx, ICA512 Ab+ (1.3 (nl<1.0)) 1 yr after dx but negative at dx
7. Unaffected daughter (9)
  a. Asthma

#### Genetic Testing to Identify *IL2RA* Mutations

Initial genetic testing of the proband using an in-house targeted next-generation sequencing multi-gene panel of over 40 genes known to be involved in monogenic forms of diabetes was negative. Subsequent exome sequencing in the trio pf proband and parents revealed the causative compound heterozygous mutations in the IL2RA gene. Two siblings carry only one mutation, but the other two with both mutations have evidence for autoimmunity: an older male sibling was found (at 4 or 5 years of age) to have positive diabetes autoantibodies in the absence of hyperglycemia and an older female sibling was diagnosed with autoimmune mediated pancytopenia at age 11 years IL2RA expression was markedly reduced in the three compound heterozygous children.

#### Clinical Phenotyping of IL2RA Patients

The IL2RA-deficient children have an almost complete loss of IL2-R□ cell surface expression on T cells and therefore virtually no detectable CD3+CD4+IL2RA^hi^CD127^l0^ Tregs in their blood, whereas family relatives carrying heterozygous *IL2RA* mutation display decreased IL2RA expression on their Tregs (**Extended Data Fig. 13a, b**). However, frequencies of CD3+CD4+CD127^l0^FOXP3+ T cells in IL2RA-deficient subjects resemble those in HD and IL2RA+/-individuals, thereby suggesting that Tregs may develop in the absence of IL2RA □□□□□□□□ (**Extended Data Fig. 13c**). Using a strategy to isolate Tregs without IL2RA expression, we found that CD3+CD4+CD127^lo^CD45RO+TIGIT+ Treg-enriched cells from IL2RA-deficient subjects showed a defective ability to suppress the proliferation of responder T cells (Tresps) as compared to HD counterparts **(Extended Data Fig. 13e, f, h)**. In contrast, Tregs from relatives with a single heterozygous *IL2RA* mutation could inhibit Tresp proliferation, although with suboptimum capacity (**Extended Data Fig. 13g, h**). Hence, correcting functional IL2RA expression on the surface of FXP3+ T cells from these patients may represent a valuable approach for developing an *ex vivo* gene therapy.

